# Generation of functional posterior spinal motor neurons from hPSCs-derived human spinal cord neural progenitor cells

**DOI:** 10.1101/2022.06.26.495599

**Authors:** Jax H. Xu, Yao Yao, Fenyong Yao, Jiehui Chen, Meishi Li, Xianfa Yang, Sheng Li, Fangru Lu, Ping Hu, Shuijin He, Guangdun Peng, Naihe Jing

**Affiliations:** State Key Laboratory of Cell Biology, CAS Center for Excellence in Molecular Cell Science, Shanghai Institute of Biochemistry and Cell Biology, Chinese Academy of Sciences, Shanghai, 200031, China; School of Life Science and Technology, ShanghaiTech University, Shanghai 201210, China; Center for Cell Lineage and Development, CAS Key Laboratory of Regenerative Biology, Guangdong Provincial Key Laboratory of Stem Cell and Regenerative Medicine, GIBH-HKU Guangdong-Hong Kong Stem Cell and Regenerative Medicine Research Centre, Guangzhou Institutes of Biomedicine and Health, Chinese Academy of Sciences, Guangzhou 510530, China; Guangzhou Laboratory, Guangzhou, 510005, China; Center for Cell Lineage and Atlas, Bioland Laboratory, Guangzhou, 510005, China; Institute for Stem Cell and Regeneration, Chinese Academy of Sciences, Beijing, 100101, China; University of Chinese Academy of Sciences, Beijing, China; Xinhua Hospital affiliated to Shanghai Jiao Tong University School of Medicine, Shanghai, 20023, China

## Abstract

Spinal motor neurons deficiency results in a series of devastating disorders such as amyotrophic lateral sclerosis (ALS), spinal muscular atrophy (SMA) and spinal cord injury (SCI). These disorders are currently incurable, while human pluripotent stem cells (hPSCs)-derived spinal motor neurons are promising but suffered from low-efficiency, functional immaturity and lacks of posterior cellular identity. In this study, we have established human spinal cord neural progenitor cells (hSCNPCs) via hPSCs differentiated neuromesodermal progenitors (NMPs) and demonstrated the hSCNPCs can be continuously expanded up to 40 passages. hSCNPCs can be rapidly differentiated into posterior spinal motor neurons with high efficiency. The functional maturity has been examined in detail. Moreover, a co-culture scheme which is compatible for both neural and muscular differentiation is developed to mimic the neuromuscular junction (NMJ) formation *in vitro*. Together, these studies highlight the potential avenues for generating clinically relevant spinal motor neurons and modeling neuromuscular diseases through our defined hSCNPCs.

## INTRODUCTION

Spinal motor neurons (MNs) which directly control muscle contractions are located in the ventral horn of spinal cord and are distributed across the anterior-posterior (AP) axis of the spinal domains from cervical, thoracic, lumbar to sacral (Kanning et al., 2010; Sockanathan and Jessell, 1998). MNs loss results in a spectrum of devastating disorders such as amyotrophic lateral sclerosis (ALS) and spinal cord injury (SCI) (Clowry et al., 1991; Narasimhan, 2006; Nijssen et al., 2017). However, *in vitro* production of spinal MNs and using MNs for drug discovery and disease modeling are hindered due to the inefficient MNs differentiation or inappropriate posterior identity.

Human pluripotent stem cells (hPSCs), including human embryonic stem cells (hESCs) and human induced pluripotent stem cells (hiPSCs), have the potential to generate various cell types *in vitro* and hold promise for disease modeling and cell-replacement therapies (Ebert and Svendsen, 2010; Karagiannis et al., 2019; Takahashi et al., 2007; Thomson et al., 1998; Yu et al., 2007). Many motor neuron differentiation protocols have been developed over the past decades (Ben-Shushan et al., 2015; Cutarelli et al., 2021; Du et al., 2015; Hudish et al., 2020; Lee et al., 2007; Li et al., 2005; Li et al., 2008; Patani et al., 2011; Qu et al., 2014). Generally, hPSCs were firstly induced to form neuroepithelial (NE) cells at embryonic body (EB) or monolayer culture, subsequently specified to OLIG2 positive motor neuron progenitors (MNPs) under the activation of retinoic acid (RA) and sonic hedgehog (SHH). These procedures require 3-4 weeks to differentiate hPSCs into MNPs and another 1-2 weeks to differentiate MNPs into immature MNs, in total 8-10 weeks to generate mature MNs from hPSCs. Alternatively, hPSCs and human fibroblasts can be converted into MNs within 2-3 weeks by forcing the expression of MN inducing transcription factors (Hester et al., 2011; Son et al., 2011). Of note, these inductions of motor neurons from hPSCs predominantly produce cells of mixed hindbrain/cervical axial identities, but are inept in generating high numbers of more posterior thoracic/lumbosacral spinal cord motor neurons, which are in most cases of key targets of disease and injury.

Spinal motor neuron development initiates from gastrulation, then epiblast cells under dramatic proliferation, differentiation and migration to form three germ layers along the AP axis (Lu et al., 2001; Solnica-Krezel, 2005; Tam and Behringer, 1997). However, the mechanism of AP patterning during the development of the central nervous system (CNS) remains obscure. *In vivo* transplantation experiments of chick embryos revealed that the nodes exhibited disparate competency at different embryonic stages for posterior nervous system induction (Storey et al., 1992). Besides, evidence from mutant T-box gene *Tbx6* in mouse embryos showed that *Tbx6* absent cells switched their cell fates from posterior somites into neuronal cells and formed three neural tube-like structures (Chapman and Papaioannou, 1998). Cell lineage segregation study by clonal analysis in early mouse embryos further indicated that trunk neural ectoderm and neighboring mesoderm derivatives share a common progenitor, namely neuromesodermal progenitor (NMP) (Tzouanacou et al., 2009). NMPs are localized in the caudal lateral epiblasts (CLE) of E8.5 mouse embryos, and express both *Sox2* and *Brachyury (T)* (Henrique et al., 2015). Based on these developmental trajectories, mouse and human NMPs were generated from mESCs and hPSCs and can be differentiated into neural and mesodermal cells *in vitro* (Davis-Dusenbery et al., 2014; Gouti et al., 2014; Turner et al., 2014; Wind et al., 2021). The chromatin accessibility assay showed that the regionalization mechanism along the AP axis during neural induction supports the dual origin of CNS, i.e., the brain and posterior spinal cord are generated from distinct progenitor populations (Metzis et al., 2018). Methods have been established to derive spinal motor neurons through NMPs. For example, Verrier and colleagues established a protocol to rapidly and reproducibly induce human spinal cord progenitors from NMPs-like cells, but these cells and their derivatives only expressed anterior spinal cord markers such as *HOXB4*, *HOXC6* and *HOXA7* (Verrier et al., 2018). Moreover, transplantation of spinal cord neural stem cells derived from hPSCs through NMP-stage showed functional recovery in SCI rat models (Kumamaru et al., 2018). Nevertheless, only about 50% of these NMP-like cells expressed both SOX2 and Brachyury (T). Thus, the establishment of high-purity human spinal cord neural progenitor cells (hSCNPCs) that are readily attainable for efficient differentiation of posterior spinal motor neurons remains unresolved.

In this study, we established a protocol to generate hSCNPCs from hPSCs through high-purity human NMPs. hSCNPCs showed molecular properties of spinal cord and were readily passaged up to 40 times *in vitro*. These hSCNPCs could be further differentiated into homogeneous spinal motor neurons, and exhibited posterior spinal cord identities. The spinal motor neurons after quick functional maturation could form neuromuscular junction (NMJ) like structures when co-cultured with muscle fibers.

## RESULTS

### Generation and characterization of human spinal cord neural progenitor cells from hPSCs

To generate NPCs with spinal cord property and posterior regional identity from hPSCs, we mimicked the posterior spinal cord development *in vitro* by differentiating hPSCs into hSCNPCs through high-purity human NMPs (Figure 1A). Feeder-free cultured hESCs were dissociated into single cells, seeded at 40,000 cells/cm^2^ and maintained in mTeSR medium for one day to form small clusters, then differentiated in N2B27 medium containing dual-SMAD inhibitors, FGF2 and CHIR99021 for another 3 days. High-purity human NMPs with 90% efficiency were obtained at day 4 (D4) by detecting SOX2 and Brachyury (T) double positive cells (Figures 1B and 1C). CDX2, another NMP marker was also highly expressed in hESCs-derived human NMPs (Figure S1A). To further investigate the molecular profiles of human NMPs, we performed bulk RNA sequencing on samples collected on each day from D0 to D4 and compared our NMPs with previously reported human NMPs, which were isolated by NKX1-2::GFP cell sorting (Verrier et al., 2018). Principal component analysis (PCA) demonstrated a clear differentiation trajectory from hESCs to human NMPs, and showed a high correlation between the D4-hNMPs with NKX1-2 enriched human NMPs (Figure 1D), suggesting that this protocol represents a new route to NMP generation.

**Figure 1.**
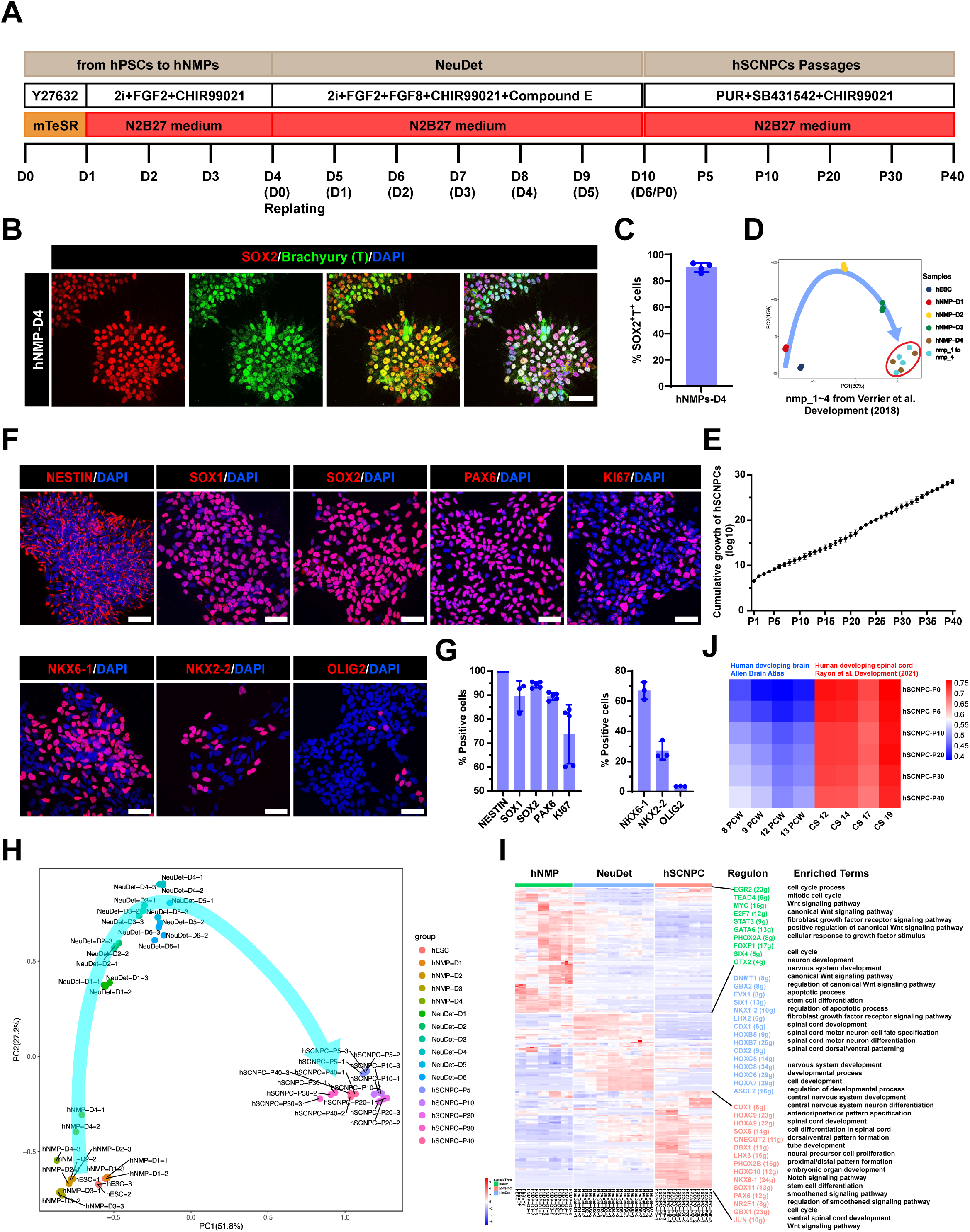
Generation and characterization of human spinal cord neural progenitor cells from hPSCs. (A) Schematic procedure for generating hSCNPCs and hiSCNPCs from hPSCs (H9, DC60-3, DC87-3) with all stages, media and factors. (B) Immunostaining of hNMPs markers SOX2 and Brachyury (T) at day 4. Scale bar, 50 μm. (C) Quantification of SOX2 and Brachyury (T) double positive cells over DAPI of hNMPs-D4. n = 4 independent experiments. Data are represented as mean ± SD. (D) Principal component analysis of time course samples during hNMPs differentiation and the previously published samples of NKX1-2-GFP sorted human nmps (Verrier et al., 2018). (E) Cumulative growth curve of hSCNPCs counts over 40 passages. n = 2 independent experiments. Data are represented as mean ± SD. (F) Immunostaining of hSCNPCs (top panel exhibited pan-NPC markers and proliferation marker; bottom panel exhibited spinal cord specific NPC markers). Scale bar, 50 μm. (G) Quantification of results shown in Figure 1F. n = 5 independent experiments. Data are represented as mean ± SD. (H) Principal component analysis of samples from hESCs (H9) to hSCNPCs at each time point shown in Figure 1A. (I) The heat-map showing three regulon groups in samples of hESCs (H9) to hSCNPCs with listing representative regulon transcription factors (numbers of predicted target genes by SCENIC in the brackets) and enriched GO terms for each regulon group. (J) Comparative analysis of whole transcriptome of hSCNPCs at different passages to dataset from Allen Brain Atlas of developing human brain and previous published dataset of developing human spinal cord.

Next, high-purity human NMPs were subjected to hSCNPCs differentiation. At D4, human NMPs were dissociated into single cells and replated with a seeding density of 100,000 cells/cm^2^ in neural determination medium (NeuDet), which contains dual-SMAD inhibitors, FGF2, FGF8, CHIR99021 and Compound E for another 6 days (Figure 1A). Real-time quantitative PCR (RT-qPCR) analysis showed that the pluripotent gene *POU5F1* and human NMP marker gene *NKX1-2* were downregulated, while *CDX2* and neural progenitor marker genes *SOX2*, *PAX6* and *NKX6-1* and HOX family genes of posterior identity such as *HOXC9* and *HOXC10* were upregulated (Figure S1D). By day 10 (D10), these hSCNPCs can be passaged in medium containing Purmorphamine, SB431542 and CHIR99021 over 40 times *in vitro* with normal karyotype (Figures 1A, 1E and S1E). Immunofluorescent staining showed that hSCNPCs expressed high percentage of pan-NPC markers NESTIN, SOX1, SOX2 and PAX6 (Figures 1F and 1G), as well as expressed moderate level of proliferation marker KI67 and spinal cord specific markers, such as NKX6-1 (∼60%), NKX2-2 (∼30%) and OLIG2 (∼5%) (Figures 1F and 1G). Human iPSCs can also be differentiated into hiSCNPCs through hiNMPs with the same protocol, indicating the robustness of this method (Figures S1B, S1C and S1F).

To further determine the dynamic changes from hESCs to hSCNPCs at the transcriptomic level, we performed bulk RNA sequencing during NeuDet stage and hSCNPCs at different passages. PCA analysis based on regulon expression showed that three major cell groups, hNMPs, differentiating neural progenitors and passaged hSCNPCs, were aligned in a timely order (Figure 1H). Regulon analysis showed that hNMPs mainly enriched with Gene Ontology (GO) terms of cell cycle process, Wnt signaling pathway and fibroblast growth factor receptor signaling pathway; NeuDet stage differentiating neural progenitors mainly enriched with stem cell differentiation and spinal cord development; passaged hSCNPCs enriched mainly with anterior/posterior pattern specification and neural precursor cell proliferation (Figure 1I). Interestingly, we observed a caudalized shift of HOX genes expression from *HOXC6-8* at NeuDet stage to *HOXC9-10* in passaged hSCNPCs (Figure 1I), suggesting an enhanced posterior signature during the derivation process. Comparison of the transcriptome of our hSCNPCs with that of human developing brain (Allen Brain Atlas) and human developing spinal cord (Rayon et al., 2021) confirmed that hSCNPCs are resembling to human developing spinal cord (Figure 1J).

Taken together, these results suggest that we have successfully obtained a neural progenitor cell line with human spinal cord features.

### Fast and direct differentiation of posterior spinal motor neurons from hSCNPCs

To test the motor neuron differentiation ability of hSCNPCs, we dissociated hSCNPCs into single cells and seeded with a density of 100,000 cell/cm^2^ on PDL and Matrigel-coated plates and differentiated in N2B27 medium containing Purmorphamine, Compound E, brain-derived neurotrophic factor (BDNF), glial cell line-derived neurotrophic factor (GDNF) and Neurotrophin-3 (NT-3) (Figure 2A). In this medium, hSCNPCs can be quickly differentiated into immature motor neurons expressed markers of HB9 (∼90%) and ISL1 (∼70%) at day 6 (D6) (Figures 2B and 2D). By day 18 (D18), hSCNPCs-derived motor neurons expressed homogeneously mature neuronal markers, such as SMI-32 (∼90%), NEUN (∼90%) and MAP2 (∼90%) (Figures 2C and 2D). These cells also expressed motor neuron subtype-specific markers, such as choline acetyltransferase (ChAT) and vesicular acetylcholine transporter (VAChT) at high percentages (Figures 2E and 2F). Other neuronal subtypes, such as glutamatergic neurons (VGLUT), GABAergic neurons (GAD67) and dopaminergic neurons (TH), were barely detectable (Figures 2E and 2F), suggesting that this method is a highly efficient spinal motor neuron differentiation protocol.

**Figure 2.**
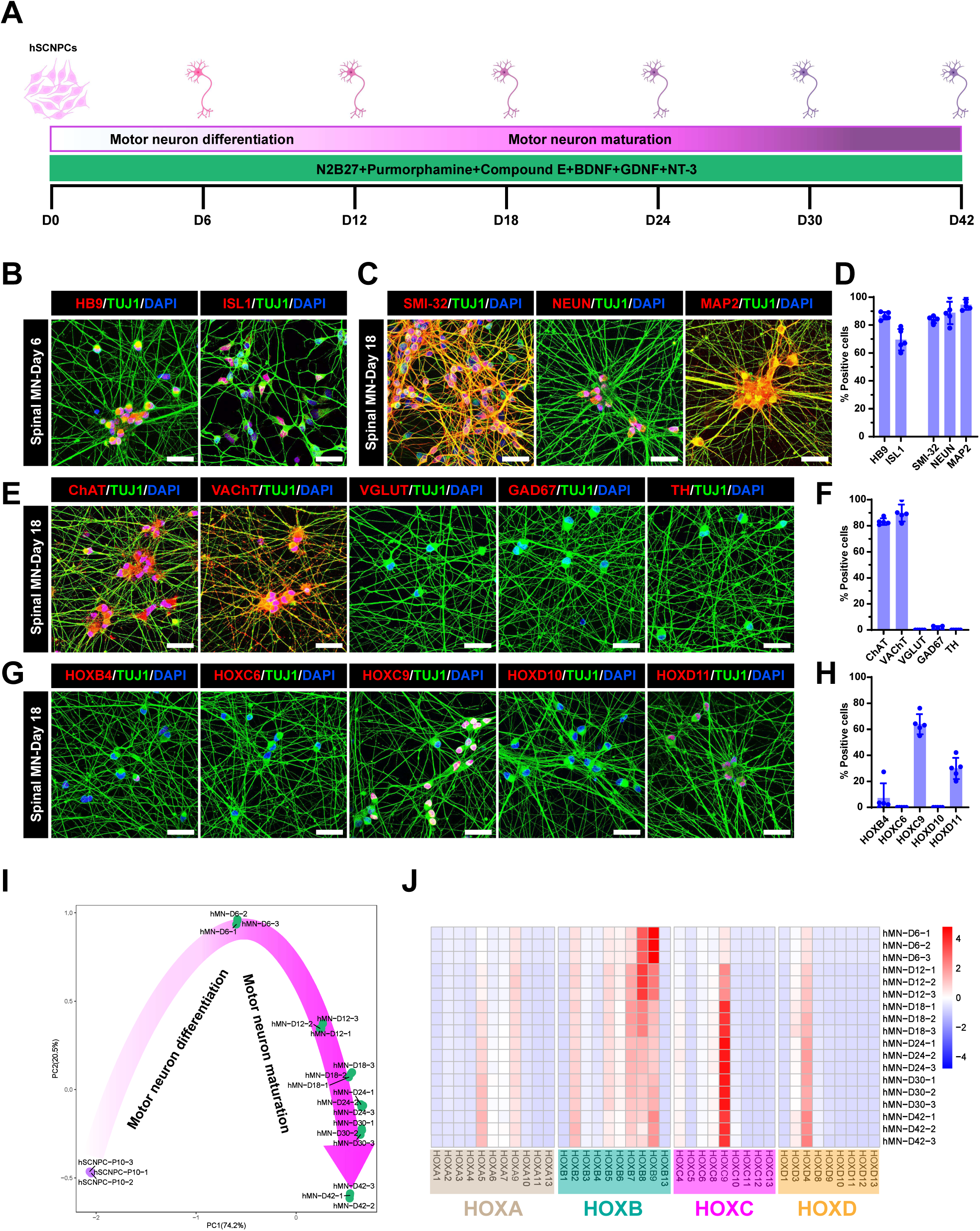
Fast and high-efficient posterior spinal cord motor neuron differentiation from hSCNPCs. (A) Schematic illustration of direct spinal motor neurons differentiation from hSCNPCs and hiSCNPCs (derived from H9, DC60-3 and DC87-3). (B) Immunofluorescent images demonstrating the expression of motor neuron markers HB9, ISL1 of hSCNPCs direct differentiated spinal motor neurons at day 6. Scale bar, 50 μm. (C) Immunofluorescent images demonstrating the expression of mature motor neuron marker SMI-32 and mature neuronal markers NEUN and MAP2 of hSCNPCs direct differentiated spinal motor neurons at day 18. Scale bar, 50 μm. (D) Quantification of results shown in Figures 2B and 2C. n = 5 independent experiments. Data are represented as mean ± SD. (E) Immunofluorescent images of neurotransmitter markers ChAT, VAChT, VGLUT, GAD67 and TH of hSCNPCs-derived spinal motor neurons at day 18. Scale bar, 50 μm. (F) Quantification of results shown in Figure 2E. n = 5 independent experiments. Data are represented as mean ± SD. (G) Immunofluorescent staining of HOX family genes HOXB4, HOXC6, HOXC9, HOXD10 and HOXD11 of hSCNPCs-derived spinal motor neurons at day 18. Scale bar, 50 μm. (H) Quantification of results shown in Figure 2F. n = 5 independent experiments. Data are represented as mean ± SD. (I) Principal component analysis of samples from hSCNPCs to spinal motor neurons at each time point shown in Figure 2A. (J) 39 HOX genes expression profiles during spinal motor neuron differentiation from hSCNPCs.

HOX genes play an important role in the rostrocaudal axis and spinal cord patterning, immunostaining analysis showed that hSCNPCs-derived spinal motor neurons mainly expressed HOXC9 (∼70%) and HOXD11 (∼30%), which were associated with thoracic and lumbar spinal domains, respectively (Figures 2G and 2H). The regional identities of hindbrain (HOXB4), cervical spinal (HOXC6) and lumbar spinal cord (HOXD10) were hardly detectable (Figures 2G and 2H). To further illustrate the expression patterns of *HOX* family genes during spinal motor neuron differentiation, we profiled 4 *HOX* clusters of 39 genes in the RNA-seq data. In line with the immunostaining results, spinal motor neurons generated from hSCNPCs specifically expressed *HOXC9* (Figure 2J). Together, these results suggest that the motor neurons this protocol induced are the posterior spinal motor neurons.

Similarly, the motor neurons derived from hiPSCs through hiSCNPCs using the same protocol also showed comparable expression levels of spinal motor neuron markers and HOX proteins (Figure S2B).

To gain insights into the transcriptomic alterations during spinal motor neuron differentiation, we employed time course RNA-seq analysis every 6 days from D0 to D30, as well as D42. The SCENIC algorithm identified 252 regulons (Figure S2C) and the PCA analysis showed a turning point at D6 (Figure 2I). Heatmap and GO term analysis showed that D0-D6 regulons were enriched with terms of spinal cord patterning, cell cycle DNA replication and neural precursor cell proliferation, and regulons of late stage were corresponding to neuron migration, neural tube formation, neuromuscular junction development and neurotransmitter secretion (Figure S2C). Hence, the spinal motor neuron differentiation process could be divided into two major events: motor neuron differentiation and maturation. We observed temporal activation of transcription factors that may account for the cell fate transition of hSCNPCs, for example, *JUN*, *GBX2*, *FOXP2*, *ONCUT3* and *SIX3* (Figure S2C). We also confirmed by RT-qPCR that the expression of NPC marker genes *NESTIN* and *NKX2-2* were downregulated, while motor neuron progenitor specific marker *NKX6-1* increased from D0 to D18 and remained the expression level thereafter, indicating that *NKX6-1* may play an important role during motor neuron fate determination and maintenance; the expression of immature neuronal marker *TUJ1* increased sharply from D0 to D6, then dramatically decreased and maintained a relatively low level; and immature motor neuron markers *HB9* and *ISL1* increased from D0 to D12 then slowly decreased (Figure S2A). In contrast, the expression of mature neuronal marker *NEUN* and mature motor neuron markers *ChAT* and *VAChT* increased from D0 to D12, and were maintained afterwards (Figure S2A).

To test whether hSCNPCs have the potential to give rise to other cell types of the spinal cord, we applied a spontaneous differentiation system to hSCNPCs. hSCNPCs were plated as single cells and differentiated in N2B27 medium without additional patterning factors. In this condition, hSCNPCs differentiated into mature neurons with centimeter-long axon bundles (Figures S2D and S2E). MicroRNA-218 (mir-218) was previously reported as an authentic marker of both rodent and human motor neurons (Amin et al., 2015; Amin et al., 2021; Hoye et al., 2018; Reichenstein et al., 2019). To compare the difference between spinal motor neurons and spontaneous differentiated neurons, time course RT-qPCR was performed for samples from both systems every 3 days from D0 to D21. The results showed that *mir-218-2* was specifically elevated in cells from spinal motor neuron differentiation system (Figure S2F). Albeit the much lower efficiency of motor neuron differentiation, we still detected HB9 positive cells in spontaneous culture. Interestingly, interneuron markers GABA and Somatostatin (SST) and astrocyte marker S100β were also presented in spontaneous differentiation (Figure S2G), indicating that hSCNPCs as neural progenitors have the capacity to generate the major neural cell lineages of spinal cord.

### Functional characterization of posterior spinal motor neuron derived from hSCNPCs

To characterize the functional maturation of the posterior spinal motor neurons, we performed whole-cell patch-clamp and CMOS-based high-density microelectrode array (HD-MEA) recordings. Spinal motor neurons co-cultured with astrocytes exhibited a mature morphology with abundant dendrites and long axons, and expressed mature motor neuron marker SMI-32 (Figures S3A and S3B). The action potentials (APs) were examined during 42 days of spinal motor neuron differentiation by whole-cell patch-clamp (Figure 3A). In response to a series of step current injections, 20% neurons could quickly generate single spike at D6 of differentiation and the mean amplitude was 24.12 ± 6.39 mV. After 12 days of differentiation, 50% neurons fired trains of APs and the mean amplitude was 41.27 ± 3.53 mV. At day 18, near 80% neurons fired repetitive APs at a mean amplitude of 47.76 ± 3.27 mV. As maturation progressively increased, 100% of recorded neurons showed sharp and repetitive spikes at 24 days, 30 days and 42 days of differentiation with mean amplitudes of 67.77 ± 2.69 mV, 71.91 ± 2.52 mV and 69.14 ± 2.63 mV, respectively (Figures 3A-3C and S3C). Statistical analysis of AP thresholds and resting membrane potentials (RMPs) showed that the mean values of AP thresholds and RMPs became more negative as the *in vitro* culture proceeded and reached a plateau after 24 days of spinal motor neuron differentiation at -32.15 ± 1.14 mV and -48.73 ± 1.12 mV, respectively (Figures 3D and 3E). A gradual reduction from ∼1 GΩ to ∼0.5 GΩ of input resistance (Rin) was also observed during spinal motor neuron maturation (Figure 3F). The F-I curves of spinal motor neurons from D6 to D42 exhibited a gradual increase in spike frequency under the increasing current injections from 0 pA to 30 pA and reached a plateau under the current injection over 30 pA (Figure 3G). Together, whole-cell patch-clamp recordings reveal that hSCNPCs-derived spinal motor neurons exhibit a fast maturation property at 24 days of differentiation, and show the highest electrophysiological function at D30 and then under a slight degeneration at D42 of differentiation.

**Figure 3.**
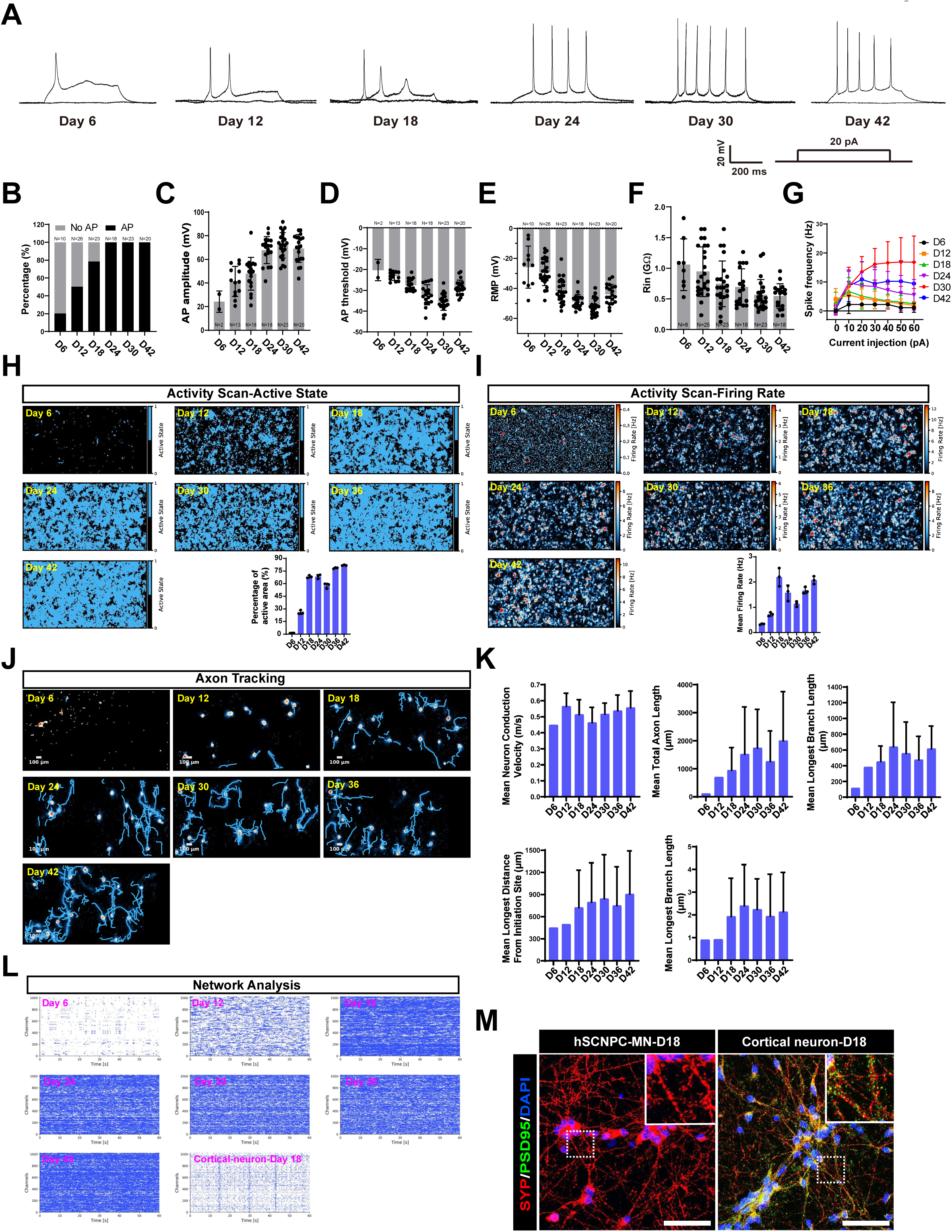
Functional characterization of hSCNPCs direct differentiated posterior spinal motor neurons. (A) Representative recording traces of hSCNPCs-derived spinal motor neurons from day 6 to day 42 in response to step current injection. (B) Percentages of hSCNPCs-derived spinal motor neurons from day 6 to day 42 with AP and no AP. N numbers of technical replicates were shown in figures from 3 independent experiments. Data are represented as mean ± SD. (C) Quantification of AP amplitude of hSCNPCs-derived spinal motor neurons from day 6 to day 42. N numbers of technical replicates were shown in figures from 3 independent experiments. Data are represented as mean ± SD. (D) Quantification of AP threshold during hSCNPCs spinal motor neurons differentiation from day 6 to day 42. N numbers of technical replicates were shown in figures from 3 independent experiments. Data are represented as mean ± SD. (E) Quantification of resting membrane potential (RMP) of hSCNPCs-derived spinal motor neurons from day 6 to day 42. N numbers of technical replicates were shown in figures from 3 independent experiments. Data are represented as mean ± SD. (F) Quantification of input resistance (Rin) of hSCNPCs-derived spinal motor neurons decreased over the differentiation process. N numbers of technical replicates were shown in figures from 3 independent experiments. Data are represented as mean ± SD. (G) Comparison of F-I curves from hSCNPCs-derived spinal motor neurons at day 6, day 12, day 18, day 24, day30 and day 42. Data were collected from 3 independent experiments. Data are represented as mean ± SD. (H) Spatial distribution maps of active spikes and the quantification results during spinal motor neuron differentiation from day 6 to day 42. n = 3 independent experiments. Data are represented as mean ± SD. (I) Representative mean firing rate and the quantification results during spinal motor neuron differentiation from day 6 to day 42. n = 3 independent experiments. Data are represented as mean ± SD. (J) Representative axon tracking maps during spinal motor neuron differentiation from day 6 to day 42. (K) Quantification results of axon tracking during spinal motor neuron differentiation from day 6 to day 42. Data were collected from 3 independent experiments. Data are represented as mean ± SD. (L) Network analysis during spinal motor neuron differentiation from day 6 to day 42 and cortical neuron at day 18 as a positive control. (M) Immunostaining of pre-(SYP, Synaptophysin) and post-synaptic (PSD95) markers of hSCNPCs-MNs and cortical neurons at day 18. Scale bar, 50 μm.

To comprehensively investigate the maturity of spinal motor neurons, we utilized the CMOS-based HD-MEA platform, which is a high-throughput and high-resolution system containing 26,400 electrodes and 1,024 simultaneous recording channels (Muller et al., 2015; Sundberg et al., 2021; Yuan et al., 2020). We first scanned and analyzed the entire HD-MEA electrode array to examine spontaneous neural activities during spinal motor neuron differentiation. The percentage of active electrodes rose rapidly from D6 to D18 and then was stabilized at about 75% thereafter (Figure 3H). The presented heatmaps of firing rate over the time course differentiation from D6 to D42, and statistical analysis of mean firing rate (MRF) showed that the time span for functional maturity is from D18 to D42, which is consistent with whole-cell patch-clamp results (Figures 3C and 3I). Then, axon tracking and spike sorting assays were performed to depict the functional images of axons and neurites of single neurons during spinal motor neuron differentiation (Figure 3J). We assessed the axon tracking metrics including mean neuron conduction velocity (m/s), mean total axon length (μm), mean longest branch length (μm), mean longest distance from initiation site (μm) and mean longest latency (ms). These results showed that all of the parameters described above were steadily increased accompanying the differentiation progresses, pointing to a step-wise electrophysiological maturation of the spinal motor neurons (Figure 3K).

Neural networks or neural circuits are featured by synchronous signals which are based on connections between pre- and postsynaptic neurons. To assess the neural network properties of spinal motor neurons, we performed network analysis during differentiation. No synchronous burst was observed from D6 to D42 after differentiation, while network oscillations were demonstrated in cortical neurons differentiated from iNSCs previously generated in our lab (Zhang et al., 2019) (Figure 3L). Moreover, we also used whole-cell patch-clamp to record spontaneous postsynaptic currents (sPSC) of spinal motor neurons at D30 of differentiation and no sPSC were recorded (Figure S3D). Although the spinal motor neurons can produce robust and abundant single burst (data not shown), they were yet to form microcircuit assembly under an *in vitro* culture system. We reasoned that the spinal motor neurons derived from hSCNPCs are of high purity and were not able to form synapses without other cell types. To test this hypothesis, immunofluorescent staining of pre- and postsynaptic markers synaptophysin (SYP) and PSD95 was performed to characterize synapse formation. The results showed that hSCNPCs-derived spinal motor neurons expressed only presynaptic marker synaptophysin but not postsynaptic marker PSD95, while iNSCs differentiated cortical neurons expressed them both (Figure 3M).

Taken together, both whole-cell patch-clamp and CMOS-based HD-MEA results reveal that hSCNPCs differentiated spinal motor neurons exhibit a fast mature property from D18 onward.

### Generation and characterization of 3D spinal motor neuron spheroids from hSCNPCs

Spinal motor neurons aggregate to form motor columns and pools in the spinal cord during vertebrate development, of which the somata reside in the spinal cord while the axons extend out to target muscles (Dasen et al., 2005; De Marco Garcia and Jessell, 2008; Guthrie, 2004). In order to mimic *in vivo* spinal motor neuron aggregates and neuron-muscle interaction *in vitro*, we generated three-dimensional (3D) spinal motor neuron spheroids from hSCNPCs and tried to find the suitable conditions for co-culture spinal motor neurons with muscle cells. Single hSCNPCs suspension was prepared and seeded in V-bottom 96-well plates at 50,000 cells/well to differentiate into 3D spinal motor neuron spheroids in the presence of Purmorphamine, Compound E, BDNF, GDNF, NT-3 and CultureOne^TM^ supplement in N2B27 medium (Figure 4A). Time course RT-qPCR showed that the expression levels of spinal cord neural progenitor marker genes *NESTIN*, *NKX2-2* and immature motor neuron marker genes *TUJ1*, *HB9* and *ISL1* declined in this 3D differentiation system; on the contrary, the expression of mature spinal motor neuron marker genes *NEUN*, *ChAT* and *VAChT* gradually increased until D12 (Figure 4B). To further confirm the spinal motor neuron feature, maturity and regional identity of 3D spinal motor neuron spheroids at D12 of differentiation, we performed immunofluorescent staining of cryosection spheroids at D12. The results showed that hSCNPCs-derived 3D spinal motor neuron spheroids homogeneously expressed motor neuron markers HB9 and SMI-32, mature neuronal markers NEUN and MAP2 and mature motor neuron markers ChAT and VAChT (Figures 4C and 4D). For regional identity, many cells (∼60%) of spinal motor neuron spheroids expressed HOXC9, but no cell expressed HOXB4 (Figures 4E and 4F). Collectively, these results suggest that the suitable time for spinal motor neuron spheroids to co-culture with muscle cells is at differentiation D12.

**Figure 4.**
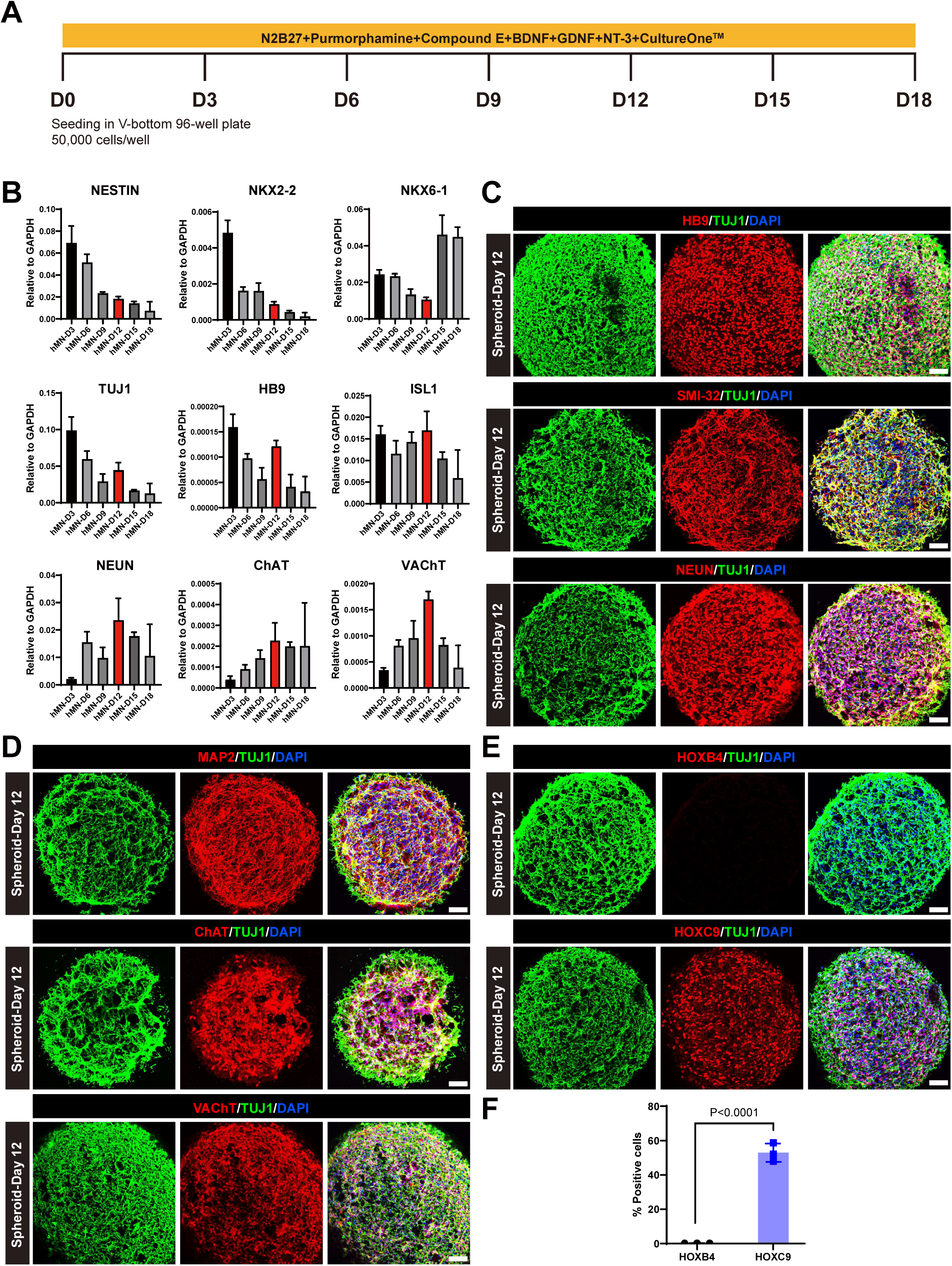
Three-dimensional spinal motor neuron spheroids differentiation from hSCNPCs. (A) Schematic illustration of the differentiation process of three-dimensional spinal motor neuron spheroids. (B) Fold changes of marker genes expression (relative to the expression level of GAPDH) during 3D spinal motor neuron spheroids differentiation. n = 4 independent experiments. Data are represented as mean ± SD. (C) Immunofluorescent staining of HB9, SMI-32 and NEUN in spinal motor neuron spheroids at day 12 after differentiation. Scale bar, 50 μm. (D) Immunofluorescent staining of MAP2, ChAT and VAChT in spinal motor neuron spheroids at day 12 after differentiation. Scale bar, 50 μm. (E) Immunofluorescent staining of HOXB4 and HOXC9 in spinal motor neuron spheroids at day 12 after differentiation. Scale bar, 50 μm. (F) Statistic analysis and quantification of Figure 4E. n = 3 independent experiments. Data are represented as mean ± SD. Student’s t test. P value was presented in the figure.

### Formation pattern of complex and pretzel-shaped acetylcholine receptor aggregates during C2C12 myogenic differentiation

One of the most important roles of spinal motor neurons is to form neuromuscular junctions (NMJs) with targeted muscles, leading to muscle innervation. To model NMJ formation *in vitro* with hSCNPCs-derived spinal motor neurons, a robust and reproducible co-culture system of spinal motor neurons and skeletal muscles is a prerequisite. To obtain an optimized myotube differentiation condition, we tested four combinations of different cell seeding densities and durations of propagation stage, then differentiated the cells for another 6 days (Figure S4A). Immunofluorescent staining of mature muscle fiber marker myosin heavy chain (MyHC) and fluorescent α-bungarotoxin (α-BTX) staining showed that C2C12 cells differentiated into the most complex structure and the highest numbers of AChR clusters with an initial seeding density of 40,000 cells/cm^2^ and 2 days propagation (Figures S4B and S4C). Thus, we established a differentiation protocol to proliferate C2C12 cells in propagation medium containing 10% FBS-DMEM for 2 days and subjected these cells to differentiate for 14 days with 2% horse serum (HS) (Figure 5A). Immunostaining analysis showed that AChR accumulated into small plaques at day 4 of myogenic differentiation, which became branched forms and pretzel-like shaped clusters at day 6, and then diffused along the myotubes at day 8 (Figures 5B and S4C). We also quantified the fiber length, fiber width, nuclei numbers per fiber, MyHC surface area, AChR area and the ratio of AChR to MyHC. The results showed that C2C12 fused into myotube and gradually matured to form muscle fibers from day 0 to day 6, and then degenerated afterwards (Figure 5C).

**Figure 5.**
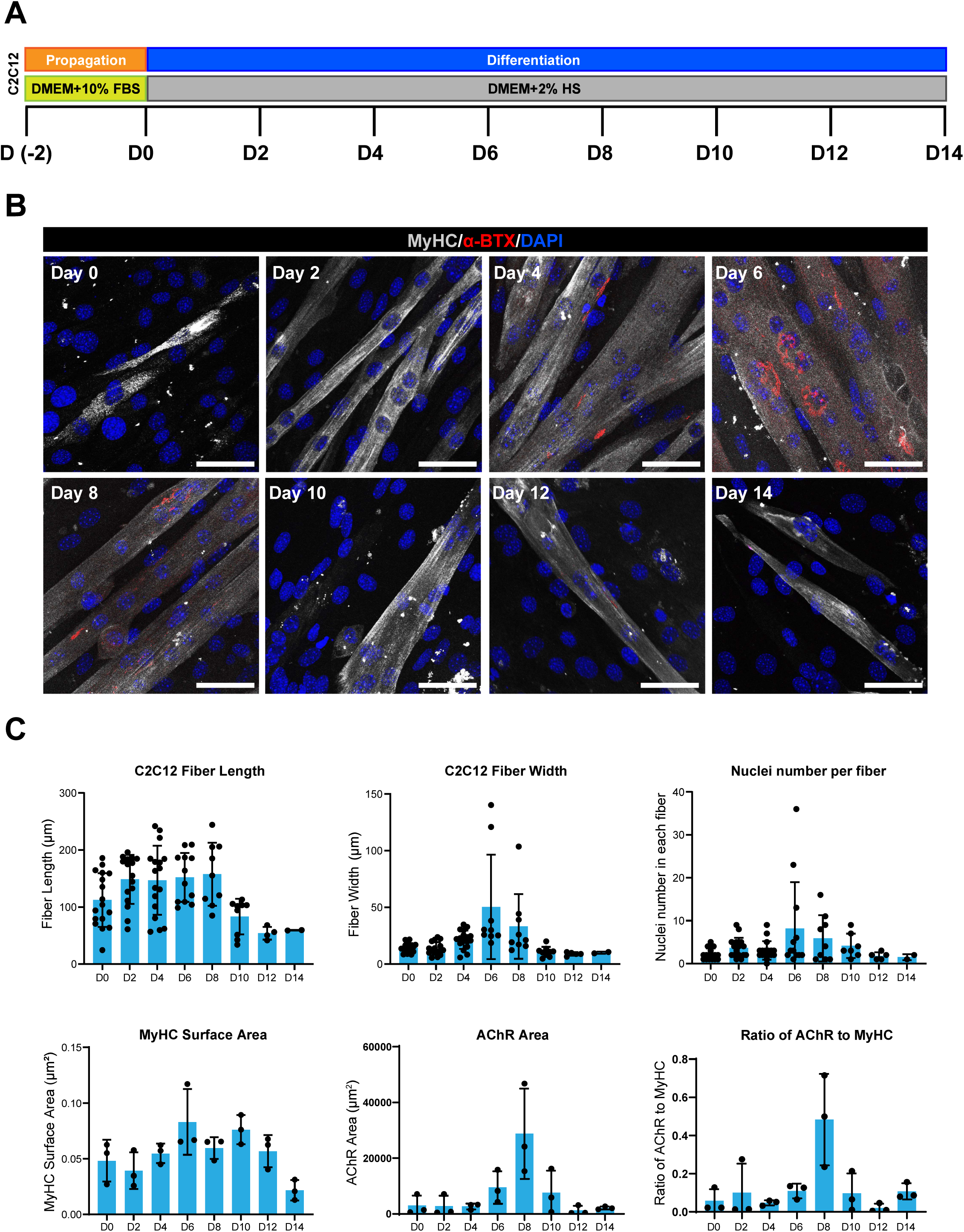
Step-wise myogenic differentiation of C2C12. (A) Schematic procedure of C2C12 differentiation. (B) Immunostaining of muscle fiber marker (MyHC) and AChR during muscle fiber differentiation of C2C12 from day 0 to day 14. Scale bar, 50 μm. (C) Quantification results of Figure 5B. Data were collected from 3 independent experiments. Data are represented as mean ± SD.

Taken together, we have established a protocol to differentiate C2C12 into mature muscle fibers with abundant and pretzel-shaped aneural AChR clusters.

### Co-culture spinal motor neuron spheroids and C2C12-derived muscle fibers to form NMJ-like structures *in vitro*

After obtaining the proper differentiation conditions for muscle cells and spinal motor neuron spheroids, we next sought to find a suitable condition to co-culture these two types of cells to model the NMJ formation *in vitro*. To this end, we first attempted to determine the co-culture medium by testing different combinations of basal medium (DMEM or N2B27), chemical additives (with or without supplementation of Purmorphamine and Compound E (P/E)) and serum (horse serum (HS) or Ultroser^TM^ G serum substitute (UG)). Having separately cultured these two types of cells in these different co-culture media for 2 days, we found that 0.2% UG and DMEM medium supported both 3D spinal motor neuron spheroids differentiation and C2C12 myotube fusion more efficiently than the other conditions, and Purmorphamine or Compound E showed no additive effect (Figures 6A and 6B). Furthermore, RT-qPCR confirmed that mature motor neuron marker genes *NEUN*, *VAChT* and *ChAT* and AChR associated markers *musk*, *chrng* and *chrne* were highly expressed in this condition (Figures 6C and 6D). Hence, we have obtained a co-culture medium that facilitates both motor neuron and myogenic differentiation.

**Figure 6.**
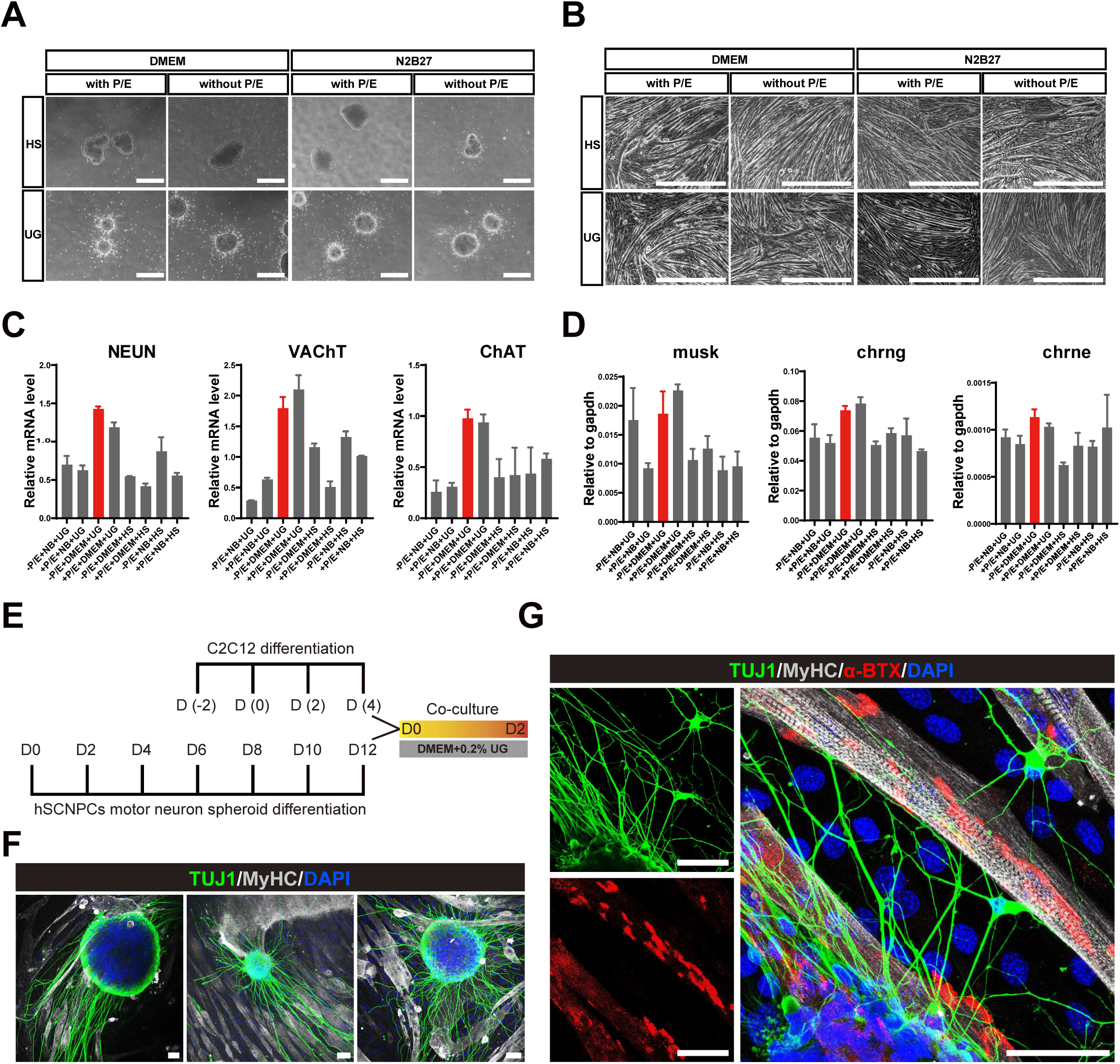
Co-culture of hSCNPCs-derived spinal motor neuron spheroids and C2C12-derived muscle fibers to form neuromuscular junctions *in vitro*. (A) Representative bright-field images of the differentiated spinal motor neuron spheroids based on differential co-culture medium compositions. Scale bar, 100 μm. (B) Representative bright-field images of the differentiated muscle fibers based on differential co-culture medium compositions. Scale bar, 100 μm. (C) Fold changes of mature motor neuron marker genes (NEUN, ChAT and VAChT) expression on differential co-culture medium. Data were collected from 2 independent experiments. Data are represented as mean ± SD. (D) Fold changes of AChR markers (musk, chrng and chrne) genes expression on differential co-culture medium. Data were collected from 2 independent experiments. Data are represented as mean ± SD. (E) Schematic illustration of co-culture model. (F) Immunofluorescent images demonstrating motor neurons extending patterns when co-cultured with muscle fibers. Scale bar, 50 μm. (G) Immunofluorescent staining of neuronal marker (TUJ1), mature muscle marker (MyHC) and AChR marker (α-BTX) on co-cultured cells. Scale bar, 50 μm.

To co-culture motor neurons and muscle cells, we added differentiation day 12 spinal motor neuron spheroids onto day 4 muscle fibers derived from C2C12 myogenic differentiation, then cultured them in the DMEM medium supplemented with 0.2% UG for 2 days (Figure 6E). Immunostaining analysis showed that spinal motor neuron spheroids extended neurites following the orientations of muscle fibers (Figure 6F), indicating that muscle fibers may have a guiding cue on axonal growth of motor neurons. Furthermore, MyHC, TUJ1 and α-BTX co-staining showed that NMJ-like structures were formed *in vitro* after co-culturing hSCNPCs-derived 3D spinal motor neuron spheroids with C2C12-derived muscle fibers for 2 days (Figure 6G).

Taken together, we have established a co-culture system, in which both spinal motor neurons and muscle cells could differentiate to maturation and form NMJ-like structures *in vitro*.

## DISCUSSION

In this study, we differentiated hPSCs into self-renewal hSCNPCs through human NMPs. These neural progenitor cells express homogeneous pan-NPC markers as well as spinal cord specific markers and can be passaged over 40 times *in vitro*. The hSCNPCs could be further differentiated into high-purity motor neurons with posterior spinal cord regional identity. The spinal motor neurons derived from hSCNPCs could be electrophysiologically mature after long-term differentiation. Moreover, co-culture of hSCNPCs-derived spinal motor neurons with C2C12-derived muscle fibers could form NMJ-like structures *in vitro*.

There is enormously regional diversity in the central nervous system. Conventionally, the generation of regional specific NPCs from hPSCs follows a general principle of neural development, hypothesized as an “activation transformation” paradigm. This model proposes that cells first undergo neural induction (“activation”) to form default anterior neural fates and caudal regions of motor neurons are generated by induction under caudalising signals such as RA and ventralizing factor SHH (“transformation”) (Mangold, 1933; Nieuwkoop and Nigtevecht, 1954; Stern, 2001). *In vitro* stepwise differentiation of motor neuron progenitors (MNPs) commonly contains two major steps. hPSCs are first differentiated into neuroepithelial (NE) cells within two weeks, then NE cells are specified to MNPs in the presence of RA and SHH (Du et al., 2015; Hu and Zhang, 2009; Lee et al., 2007; Li et al., 2005). Meanwhile, dual-SMAD inhibitors are used to increase the differentiation efficiency of NE cells and shorten the differentiation time (Chambers et al., 2009). However, the most caudal markers characterized in those studies are relative to hindbrain and cervical region of anterior spinal cord (Sances et al., 2016). Mechanistic studies also reveal that RA treatment leads to the binding of retinoic acid receptors (RARs) to *Hox1-5* chromatin domains (Mahony et al., 2011; Mazzoni et al., 2013). Increasing evidences have indicated that posterior spinal cord has a distinct developmental origin from NMPs (Gouti et al., 2014; Metzis et al., 2018; Tzouanacou et al., 2009). Human spinal cord neural stem cells have been generated from hPSCs through a transient stage of NMP, but the efficiency of SOX2 and Brachyury (T) double positive NMPs is only around 50% (Kumamaru et al., 2018), raising the potential safety concerns. In this study, we have optimized human NMPs differentiation system and obtained over 90% efficiency (Figure 1). Importantly, this protocol is stable and reproducible due to optimized seeding density and standard operation procedures which could facilitate its clinical usage. The hSCNPCs can be derived from human NMPs and exhibit spinal cord specific features by expressing NKX6-1, NKX2-2 and OLIG2 with continuously passage property *in vitro* (Figure 1).

As one of the vital cell types in the spinal cord, motor neurons (MNs) are of great clinical importance in spinal cord injury as well as motor neuron diseases such as ALS and SMA (Fujimori et al., 2018; Olmsted et al., 2021; Sances et al., 2016; Wang et al., 2013). Conventionally, differentiation of postmitotic motor neuron has disadvantages in time consumption, low efficiency and particularly incapable of making caudal spinal motor neurons (Hu and Zhang, 2009; Sances et al., 2016). The hSCNPCs based spinal motor neuron differentiation is robust and only takes 18 days to generate ChAT and VAChAT positive mature spinal motor neurons, and these cells express posterior spinal cord HOX markers such as HOXC9 and HOXD11 (Figure 2). Furthermore, the whole-cell patch-clamp and HD-MEA recordings showed that the hSCNPCs-derived spinal motor neurons can fire single AP as early as 6 days of differentiation and over 80% cells can fire APs at day 18. By day 30, the membrane properties of spinal motor neurons reach to the most mature function (Figure 3). HD-MEA analyses also showed a consistent maturation pattern as patch-clamp electrophysiology illustrated (Figure 3).

In motor neuron diseases, the core synapse and therapeutic relevant structure is the neuromuscular junction (NMJ) (Kariya et al., 2014; Picchiarelli et al., 2019). *In vitro* modeling of NMJ has been challenging, which demands proper juxtaposition of both spinal motor neuron and skeletal muscle differentiation (Bakooshli et al., 2019; Guo et al., 2011; Martins et al., 2020; Osaki et al., 2020). The established protocols for generating NMJ *in vitro* can hardly form pretzel-shaped NMJ structures. By optimizing the C2C12 differentiation process, we showed that the AChR cluster formation begins at day 4 and degenerates at day 8 of *in vitro* differentiation (Figure 4). Moreover, we also have established a protocol to differentiate hSCNPCs into 3D spinal motor neuron spheroids which can better mimic the positional relation between spinal motor neurons and skeletal muscle fibers (Figure 5). Spinal motor neurons and skeletal muscle fibers favor different culture conditions (Barbeau et al., 2020). For example, muscle fiber differentiation needs horse serum whereas serum does not benefit neural differentiation. Thus, we have systematically screened different co-culture conditions, and have found that DMED medium supplemented with Ultroser^TM^ G serum substitute is most suitable for co-culture of hSCNPCs-derived spinal motor neuron spheroids and C2C12-derived muscle fibers. Finally, we have obtained an efficient co-culture system for spinal motor neurons and skeletal muscles, and reproduced NMJ-like structures *in vitro* (Figure 6).

Adult mammals have limited regeneration of CNS, while stem cells-derived NPCs hold promises to restoring functional neural circuits *in vivo* (Varadarajan et al., 2022). Studies of cell transplantation therapy of SCI also provide evidences of motor neuron progenitors on the recovery of locomotor functions (Ceto et al., 2020; Kadoya et al., 2016; Kumamaru et al., 2018; Kumamaru et al., 2019). The hSCNPCs-derived from high-purity NMPs and exhibit spinal cord specific properties, and can be largely expanded to meet the need of high-volume cell transplantation. Besides, the ability to form NMJ and 3D sophistical structure would facilitate the mechanistic studies of human spinal motor neurons in development and disease. We expect the availability of hSCNPCs will hold great potential for regenerative medicine and translational study.

## ACKNOLEDGEMENTS

This work was supported in part by the National Key Basic Research and Development Program of China (2019YFA0801402, 2018YFA0800100, 2018YFA0108000, 2018YFA0107200), “Strategic Priority Research Program” of the Chinese Academy of Sciences, Grant No. (XDA16020501, XDA16020404), National Natural Science Foundation of China (32130030, 31630043, 31871456, 31900454).

## AUTHOR CONTRIBUTIONS

J.H.X. and N.J. conceived the project. J.H.X., Y.Y. and M.L performed the experiments and collected the data. J.H.X. and Y.Y. designed the experiments, analyzed the data and made figures. F.Y. performed patch-clamp electrophysiology. X.Y. produced and J.C. analyzed the RNA-seq data. J.H.X., G.P. and N.J. wrote the manuscript. N.J. supervised the study. All authors read and approved the final manuscript.

## KEY RESOURCES TABLE

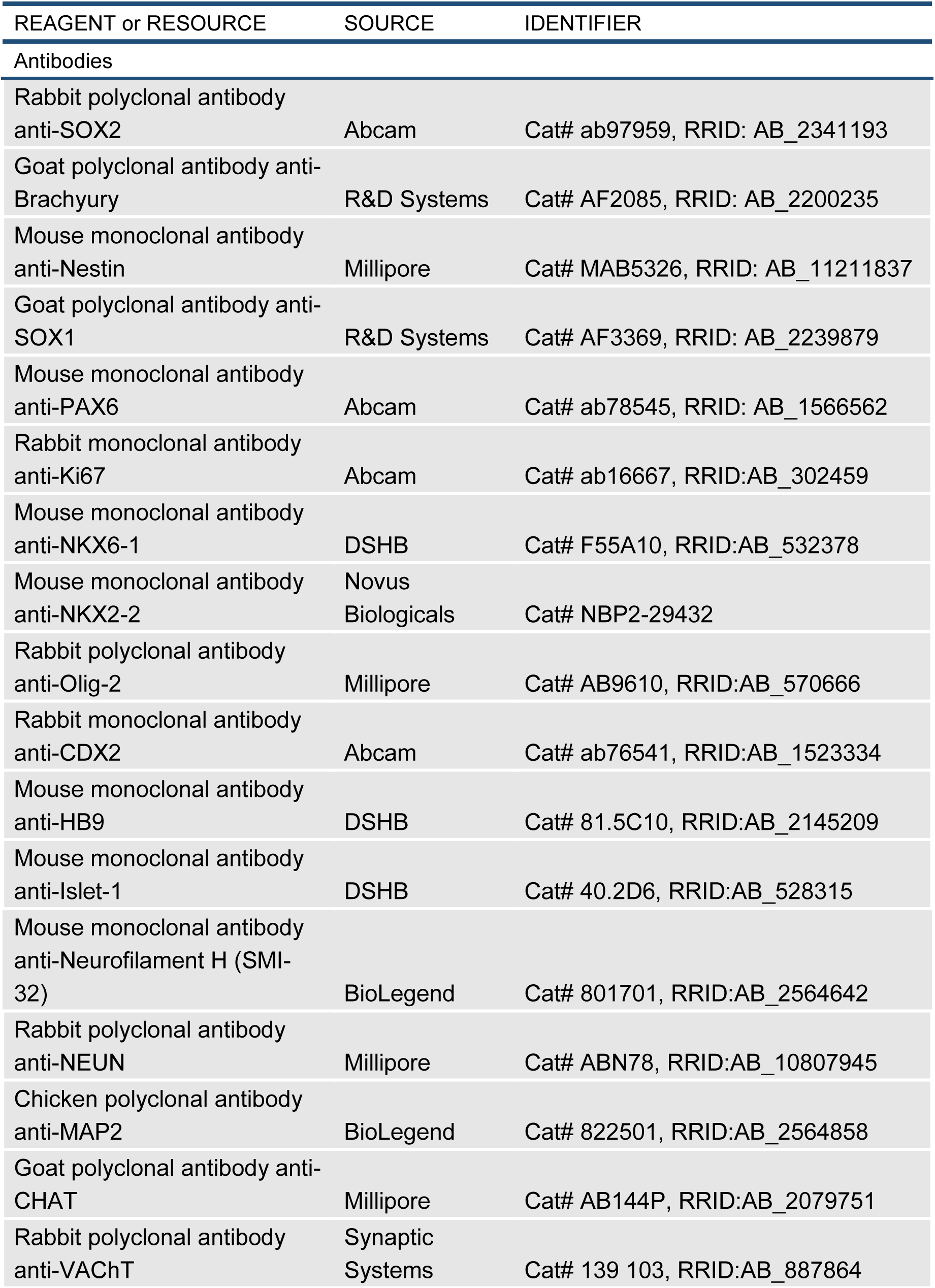
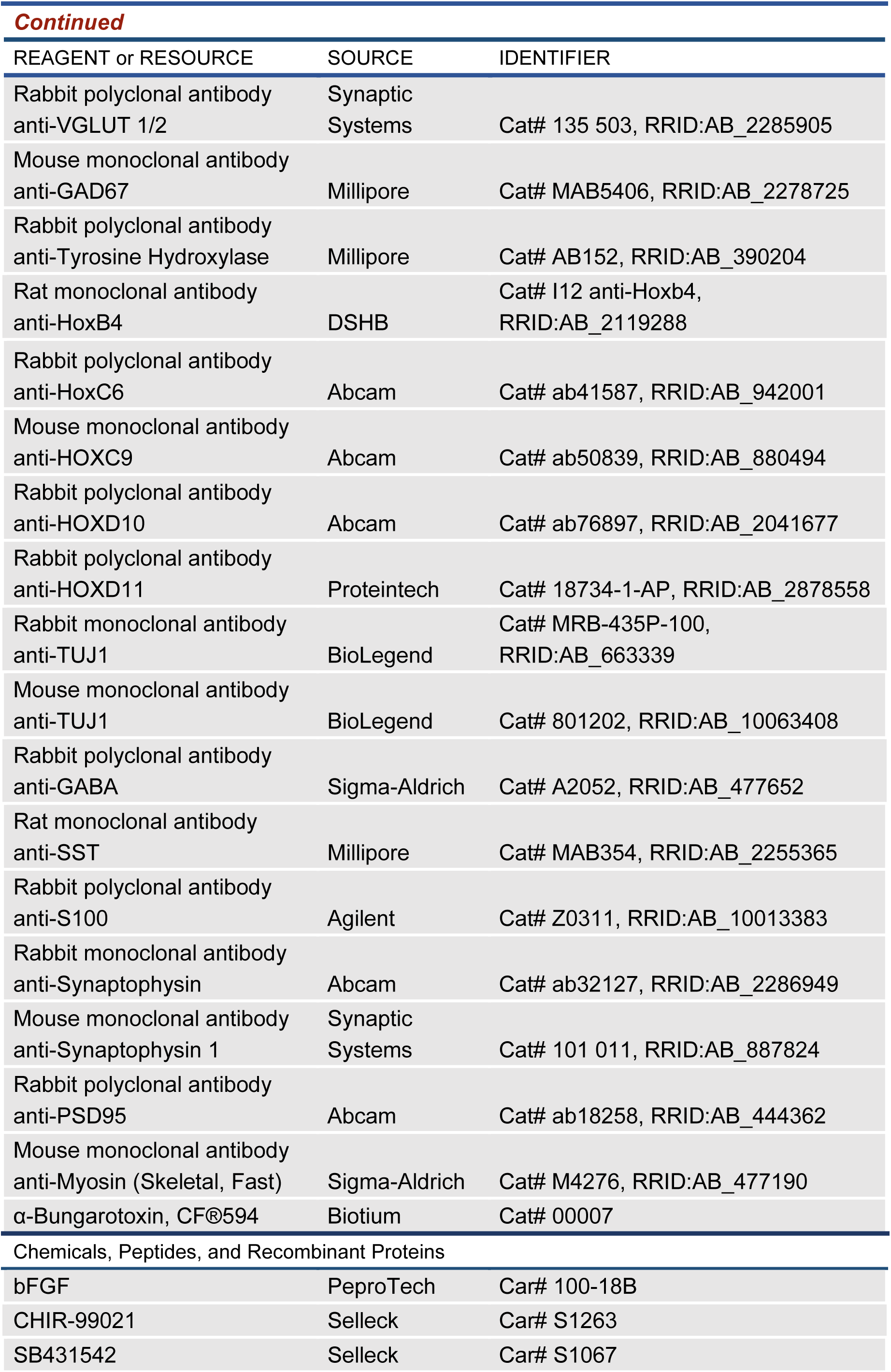
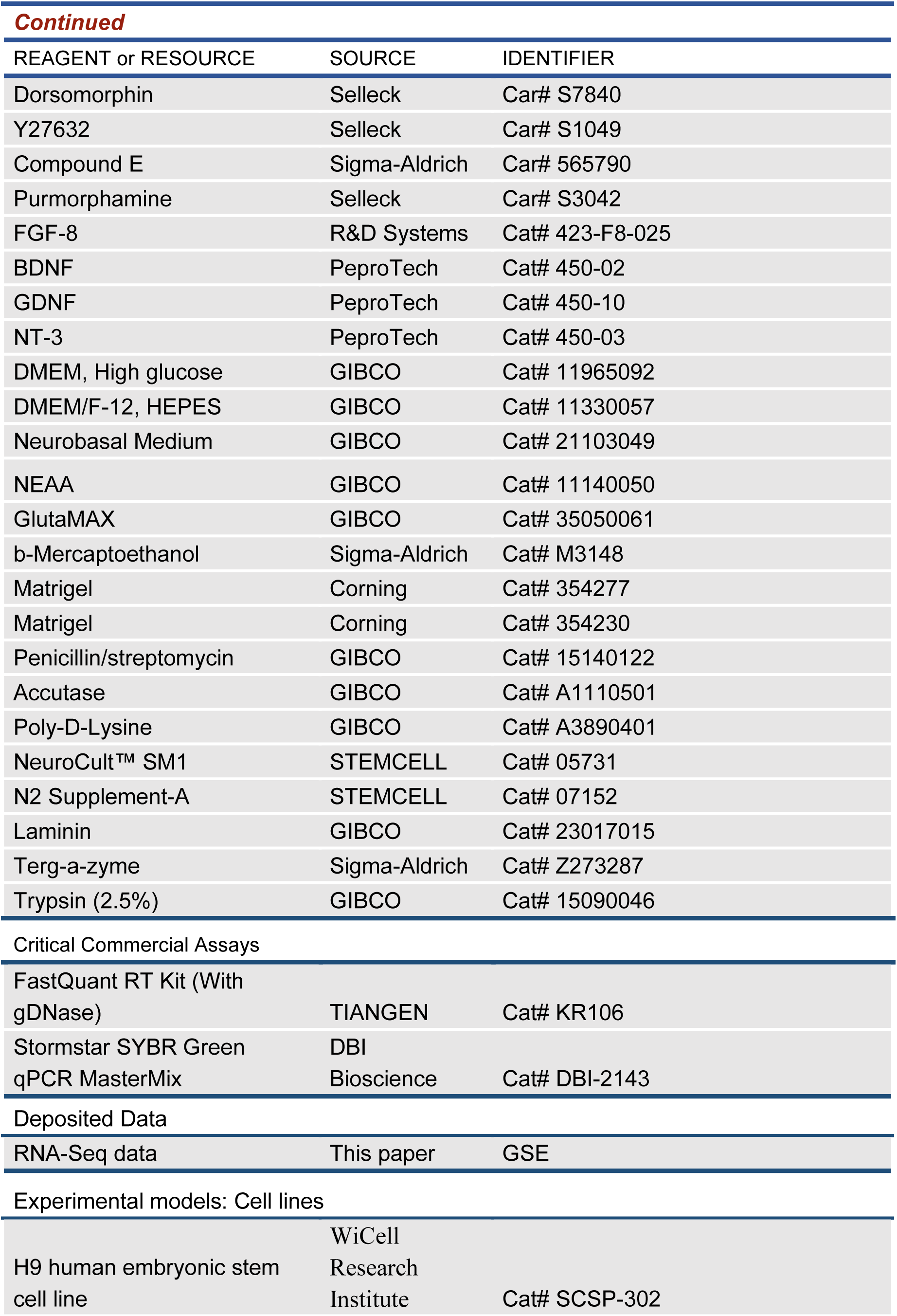
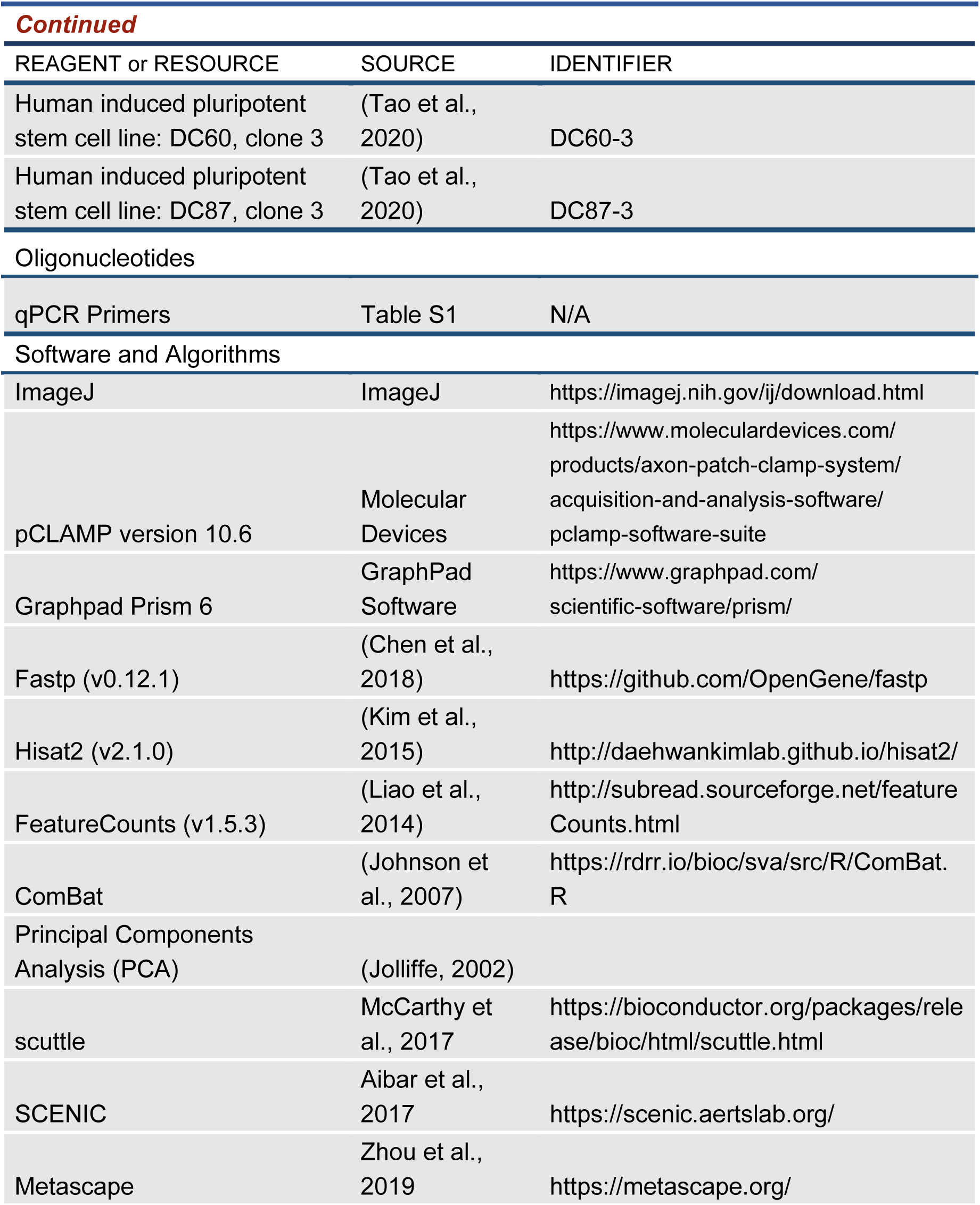

## STAR∗METHODS

### LEAD CONTACT AND MATERIALS AVAILABILITY

For further information and requests for reagents and resource, please contact the leading correspondence Naihe Jing. All materials and reagents will be made available upon installment of a material transfer agreement (MTA).

#### hPSCs Culture

Human embryonic stem cells (H9) (Thomson et al., 1998) was obtained from WiCell Research Institute. Human induced pluripotent stem cells (DC60-3, DC87-3) were generated from human peripheral blood mononuclear cells (PBMNC) in our lab previously (Tao et al., 2020). Undifferentiated hPSCs were cultured in mTeSR1^TM^ medium (STEMCELL Technologies) on Matrigel (Corning, hESCs-qualified)-coated cell culture plates. mTeSR1^TM^ medium was changed every day and hPSCs clones were disassociated into small clumps with ReLeSR (STEMCELL Technologies) as required and passaged every 3-4 days.

#### Generation of human neuromesodermal progenitors from hPSCs

hPSCs (H9, DC60-3 and DC87-3) were disassociated into single cells with Accutase (Gibco) for 6-10 min at 37 ℃ incubator. hPSCs single cell suspension were counted and plated at a density of 45,000 cells/cm^2^ in mTeSR1^TM^ medium on Matrigel (Corning)-coated cell culture plates for 1 day then cell culture medium was changed into hNMP differentiation medium at the second day and culture medium was changed every day up to day 4. 100 ml of hNMP differentiation medium composited with 47.5 ml of DMEM/F12 (Gibco), 47.5 ml of Neurobasal (Gibco), 1 ml of N2 (STEMCELL Technologies, 100X), 2 ml of NeuroCult™ SM1 Without Vitamin A (STEMCELL Technology, 50X), 1 ml of NEAA (Gibco), 1 ml of GlutaMAX (Gibco), 20 ng/ml FGF-2 (PeproTech), 3 μM CHIR99021 (Selleck), 10 μM SB431542 (Selleck), 20 μM Dorsomorphin (Selleck).

#### Generation of human spinal cord neural progenitor cells from hNMPs

hNMPs were disassociated into single cells with Accutase (Gibco), and plated at a density of 100,000 cells/cm^2^ in hSCNPCs induction medium on Matrigel-coated cell culture plates. hSCNPCs induction medium was changed every day up to day 6. 100 ml of hSCNPCs induction medium composited with 47.5 ml of DMEM/F12 (Gibco), 47.5 ml of Neurobasal (Gibco), 1 ml of N2 (STEMCELL Technologies, 100X), 2 ml of NeuroCult™ SM1 Without Vitamin A (STEMCELL Technology, 50X), 1 ml of NEAA (Gibco), 1 ml of GlutaMAX (Gibco), 100 ng/ml FGF-2 (PeproTech), 100 ng/ml FGF-8 (PeproTech), 4 μM CHIR99021 (Selleck), 10 μM SB431542 (Selleck), 20 μM Dorsomorphin (Selleck), 0.2 μM Compound E (Sigma-Aldrich).

#### hSCNPCs culture

hSCNPCs were cultured in N2B27 medium supplemented with 2 μM SB431542, 3 μM CHIR99021, 0.25 μM Purmorphamine, 1X NEAA (Gibco), 1X GlutaMAX (Gibco) and 0.1 mM 2-Mercaptoethanol (Sigma-Aldrich), on Matrigel (Corning)-coated cell culture plates. hSCNPCs were dissociated into single cells with Accutase (Gibco), and were plated on Matrigel (Corning)-coated plates in a density of 300,000/cm^2^ in hSCNPCs maintenance medium.

#### Spinal motor neuron differentiation from hSCNPCs

To differentiate hSCNPCs into motor neurons, hSCNPCs were dissociated into single cells with Accutase, counted and plated at a density of 100,000/cm^2^ on pre-treated dish (Dish pre-treated with PDL coated for 2 hours, wash 3 times with sterilized water and then Matrigel coated for 2 hours). 100 ml of motor neuron differentiation medium composited with 47.5 ml of DMEM/F12 (Gibco), 47.5 ml of Neurobasal (Gibco), 1 ml of N2 (STEMCELL Technologies, 100X), 2 ml of NeuroCult™ SM1 Without Vitamin A (STEMCELL Technology, 50X), 1 ml of NEAA (Gibco), 1 ml of GlutaMAX (Gibco), 1 μM Purmorphamine, 0.2 μM Compound E, 10 ng/ml BDNF, 10 ng/ml GDNF, 10 ng/ml NT-3. 10 μM Y27632 was added at the plating day. The motor neuron differentiation medium was changed every other day.

#### Spontaneous differentiation of hSCNPCs

hSCNPCs were dissociated into single cells with Accutase (Gibco), and were plated at a density of 100,000/cm^2^ on pre-treated dish (Dishes were pre-treated with PDL for 2 hours, washed 3 times with sterilized water and then re-coated with Matrigel for 2 hours). 100 ml of spontaneous differentiation medium composited with 47.5 ml of DMEM/F12 (Gibco), 47.5 ml of Neurobasal (Gibco), 1 ml of N2 (STEMCELL Technologies, 100X), 2 ml of NeuroCult™ SM1 Without Vitamin A (STEMCELL Technology, 50X), 1 ml of NEAA (Gibco), 1 ml of GlutaMAX (Gibco). 10 μM Y27632 was added at the plating day. The spontaneous differentiation medium was changed every other day.

#### 3D spinal motor neuron spheroids differentiation from hSCNPCs

To generate 3D spinal motor neuron spheroids, hSCNPCs were digested into single cells with Accutase (Gibco), counted and distributed into V-bottom 96-well plate at a density of 50,000/well. 3D spinal motor neuron spheroids were differentiated in N2B27 medium contains Purmorphamine, Compound E, BDNF, GDNF, NT-3 and CultureOne^TM^ supplement (Gibco). Differentiation medium was changed every other day.

#### C2C12 maintenance and differentiation

C2C12 myoblasts were cultured in high-glucose DMEM medium with 10% FBS. The medium was changed every other day. To differentiate C2C12 into muscle fibers, C2C12 were dissociated into single cells with 0.05% trypsin, counted and plated in 45,000 cells/cm^2^ on pre-coated dishes (Dishes were pre-treated with PDL for 2 hours, washed 3 times with sterilized water and then re-coated with Laminin for 2 hours) in C2C12 maintenance medium. On the following day, the C2C12 maintenance medium was replaced with differentiation medium (DMEM with 2% horse serum). Then the differentiation medium was changed every other day.

#### Co-culture of spinal motor neuron spheroids and C2C12-derived muscle fibers

To co-culture spinal motor neurons spheroids with C2C12-derived muscle fibers, we pre-differentiated C2C12 as described above, after two days of C2C12 differentiation, spinal motor neuron spheroids pre-differentiated for 12 days were plated onto C2C12-derived muscle fibers in co-culture medium (DMEM supplemented with 0.4% Ultroser^TM^ G and 10 ng/ml BDNF and 10 ng/ml GDNF and 10 μg/ml insulin. The co-culture medium was changed every other day.

#### Electrophysiology of spinal motor neurons

Whole-cell patch-clamp recordings were performed as previously described (Zhang et al., 2020). hPSCs-derived spinal motor neurons were cultured in a 35 mm petri dish. For recording, the neurons were perfused with the ACSF solution (in mM): 126 NaCl, 4.9 KCl, 1.2 KH_2_PO_4_, 2.4 MgSO_4_, 2.5 CaCl_2_, 26 NaHCO_3_, 20 Glucose. Recording pipettes (8-10 MΩ) were fabricated using a P1000 micropipette puller (Sutter Instrument, USA) and filled with internal solution consisting of (in mM): 136 K-gluconate, 6 KCl, 1 EGTA, 2.5 Na_2_ATP, 10 HEPES. For recording action potentials, cells were recorded at a holding potential of -70 mV and a series of step depolarizing currents were injected into cells in a current-clamp mode. All the recordings were performed using an Olympus microscope (BX51WI). The data were sampled at 10 Hz and collected with a low-pass filter at 2 kHz and analyzed using the Axopatch1500B amplifier and pCLAMP10 software (Molecular Devices) and GraphPad Prism 8.0.2.

#### HD-MEA recordings and analysis of hSCNPCs-derived spinal motor neurons

To analyze the function of hSCNPCs-derived motor neurons, we used high-content multi-electrode array system (MaxOne System, MaxWell Biosystems AG, Switzerland). MaxOne chips were pre-coated with 1% Terg-a-zyme (Sigma) overnight at room temperature and washed 3 times with sterile ddH_2_O, then transfered MaxOne Chips in a beaker filled with 75% ethanol for 1 hour in a biological safety cabinet, and then washed 3 times with sterile ddH_2_O, then 500 μl PDL was added for coating MaxOne chips for 2 hours at 37 ℃, then washed 3 times with sterile ddH_2_O and re-coated MaxOne chips with Matrigel (Corning) for 2 hours at 37 ℃. hSCNPCs were dissociated into single cells, counted and plated on MaxOne chips at a density of 300,000 cells/chip in motor neuron differentiation medium. MEA data recording started after 6 days of plating and measured every 6 days till day 42.

For MEA data recording, MaxLab Live Software (v.21.1.2. MaxWell Biosystems AG, Switzerland) was used. Assays were chosen from assay gallery which contains three assays, activity scan, network and axon recording. To be able to track axons, we first recorded the whole MaxOne HD-MEA surface using the “Activity Scan Assay” module, featured in the MaxLab Live software (MaxWell Biosystems AG, Zurich, Switzerland). 29 electrode configurations, including a total of 26,400 electrodes at a distance of 17.5 μm, were used to record the spontaneous neuronal electrical activity across the entire MaxOne HD-MEA. Each electrode configuration was recorded for 60 seconds.

In a next step, axonal signals were identified with the “Axon Tracking Assay” module of the MaxLab Live software, which uses a tailored recording strategy similar to the ones described in previous publications (Bakkum et al., 2013; Bullmann et al., 2019; Radivojevic et al., 2017). First, a set of 30 target neurons was identified by selecting the positions of 30 electrodes featuring the largest spike amplitude, keeping a minimum distance of 17.5 μm between each two target neurons. For every target neuron, a 3×3 electrodes block (distance between electrodes 17.5 μm) was selected around the central electrode. After defining the 3×3 blocks, a set of sequential 180 s recordings that covered larger array areas (300×300 to 800×800 μm^2^ around the blocks) was run, where the 3×3 central blocks were always included in every recording.

The detected spikes on the 3×3 electrode blocks were then sorted using a K-Means clustering algorithm (Lewicki, 1998) based on the action potential minimum spike amplitude. Spike sorting results were used to reconstruct the spike-triggered average extracellular waveform over the recorded array area for every identified neuron. An unsupervised object-tracking algorithm was then used to detect the path of action potential signal propagation, thus identifying (1) individual axonal branches and (2) the morphology of neuronal outgrowth per cell. The axonal conduction velocity was then calculated for every detected axonal branch.

#### Immunocytochemistry

Cells were fixed with 4% paraformaldehyde (PFA, Sigma-Aldrich) for 20 minutes at room temperature then washed 3 times with PBS. Blocking with 0.3% Triton X-100 /5% fetal bovine serum in PBS for one hour at room temperature. Primary antibodies were diluted in 0.3% Triton X-100 /5% fetal bovine serum in PBS and incubated overnight at 4 ℃. Samples were washed in PBS for three times and then incubated with secondary antibodies (1:200-1:1000) in 0.3% Triton X-100 /5% fetal bovine serum in PBS for two hours. Images were taken with Leica TCS SP8 confocal laser-scanning microscope.

#### RT-qPCR analysis

Total RNA was extracted from cultured cells by using the TRIzol reagent, and cDNA was reverse-transcribed, starting from 1000 ng of total RNA with the SuperScript III First-strand cDNA synthesis kit (Invitrogen). qPCR was performed using Mastercycler RealPlex2 (Eppendorf) and Stormstar SYBR Green qPCR MasterMix (DBI Bioscience). Data were normalized for GAPDH expression. The primers sequences used for qPCR amplification were listed in Table S1.

#### RNA-Sequencing and data analysis

Cell samples at different stages were collected and mRNA-seq libraries were constructed with NEBNext® Ultra™ II Library Prep Kit for Illumina. Qualified libraries were multiplexed and sequenced on Illumina NovaSeq 6000 according to the manufacturer’s instructions (Illumina, USA). The Sequencing mode was PE150.

For data analysis, reference genome and gene annotation files were downloaded from GENECODE (hg38). Fastq files were pre-processed by Fastp (v0.12.1) (Chen et al., 2018) with default parameters. Cleaned data were then aligned to the reference genome via Hisat2 (v2.1.0) (Kim et al., 2015). FeatureCounts (v1.5.3) (Liao et al., 2014) was used to count the reads mapped to each gene.

##### Correlation to nmps

RNA-seq data in this paper were compared with nmps (Verrier et al., 2018) after removing batch effects by the R package “ComBat” (Johnson et al., 2007). Principal Components Analysis (PCA) (Jolliffe, 2002) was used to show the correlation of these samples.

#### Correlation to BRAINSPAN database and pseudo-bulk RNA-seq of human single cell spinal cord

The ‘Developmental Transcriptome Dataset’ of BRAINSPAN (https://www.brainspan.org/static/download.html) was also downloaded. We merged the brain samples of same stage and calculated Pearson correlation coefficients between the merged samples and RNA-seq data in this paper.

Pseudo-bulk RNA-seq of human spinal cord, GSE171892 (Rayon et al., 2021), was conducted through the R package “scuttle” (McCarthy et al., 2017). After that, samples at same stages were merged and Pearson correlation coefficients were calculated between the pseudo-bulk RNA-seq and RNA-seq data in this paper.

#### Inference of regulons and their activity

The SCENIC (Aibar et al., 2017) was used to infer the gene regulatory network (regulon). Binary regulon-activity matrix for all samples was used in principal components analysis. GO enrichment analysis of related target genes was carried out by Metascape (https://metascape.org/) with default parameters (Zhou et al., 2019).

### Statistical analysis

All statistical analyses were performed in GraphPad Prism software (GraphPad Prism 8.0.2). Cell counting, RT-qPCR data and electrophysiological data were presented as mean ± SD. Student’s t test (two-tailed) was performed for statistical analysis between two groups. Sample size (n) values were provided in the relevant text, figures, and figure legends. The statistical analyses were obtained from three independent experiments. Statistical significance was set at p values.

### Data availability

All RNA-seq data are available at the Gene Expression Omnibus (GEO) under accession number GSE205718.

---

**Figure S1.**
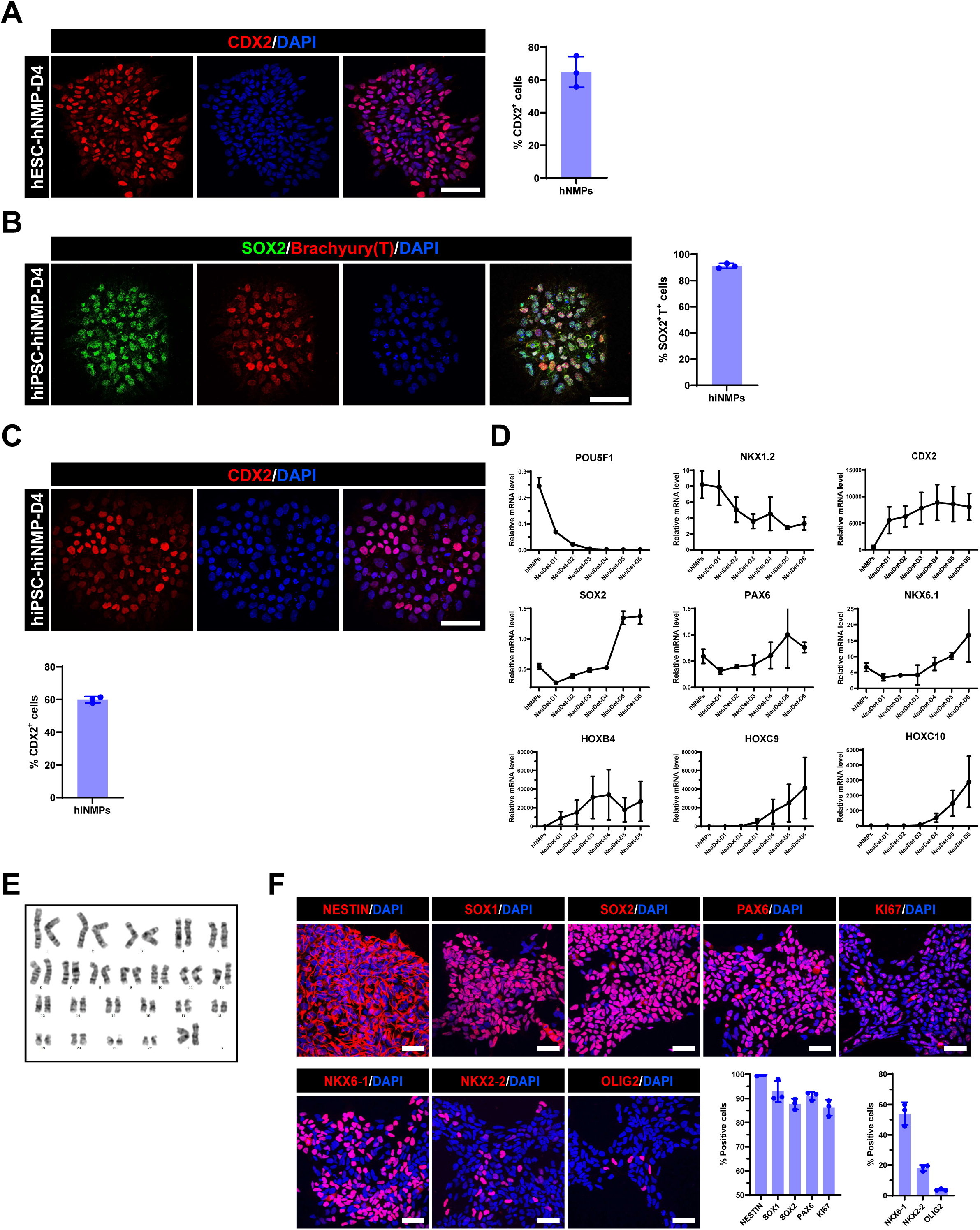
Characterization of human NMPs and hSCNPCs from hPSCs. (A) Immunofluorescent staining of human NMP marker (CDX2) and the quantification results. Scale bar, 50 μm. n = 3 independent experiments. Data are represented as mean ± SD. (B) Immunofluorescent staining of human NMP markers (SOX2 and Brachyury) of hiPSCs (DC60-3 and DC87-3)-derived NMPs and the quantification results. Scale bar, 50 μm. n = 3 independent experiments. Data are represented as mean ± SD. (C) Immunofluorescent staining of human NMP marker (CDX2) of hiPSCs (DC60-3 and DC87-3)-derived NMPs and the quantification results. Scale bar, 50 μm. n = 3 independent experiments. Data are represented as mean ± SD. (D) Relative marker genes expression of during NeuDet stage. n = 3 independent experiments. Data are represented as mean ± SD. (E) Karyotyping of hSCNPCs at P40. (F) Immunofluorescence analysis of hiPSCs (DC60-3 and DC87-3)-derived hiSCNPCs and the quantification results. Scale bar, 50 μm. n = 3 independent experiments. Data are represented as mean ± SD.

**Figure S2.**
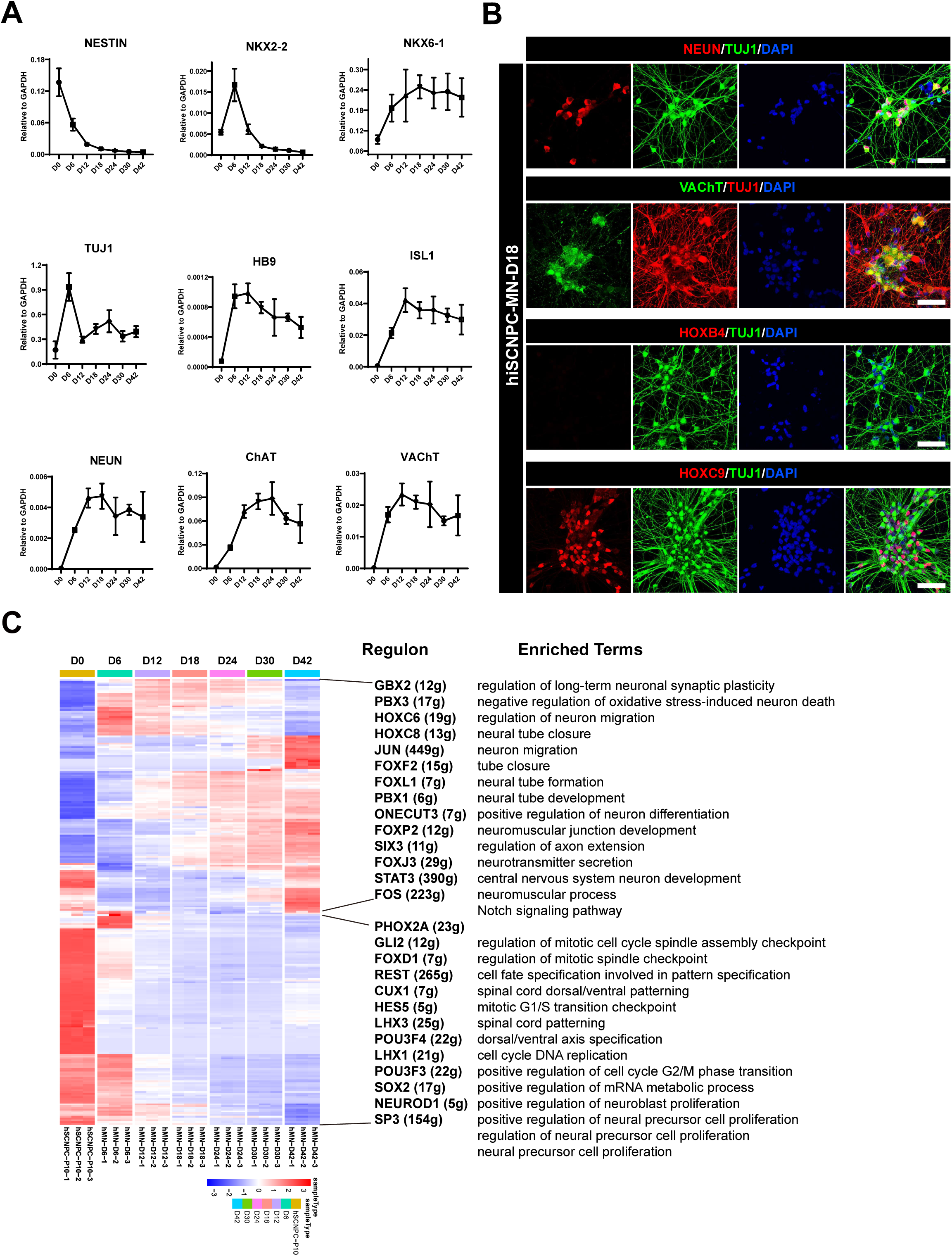
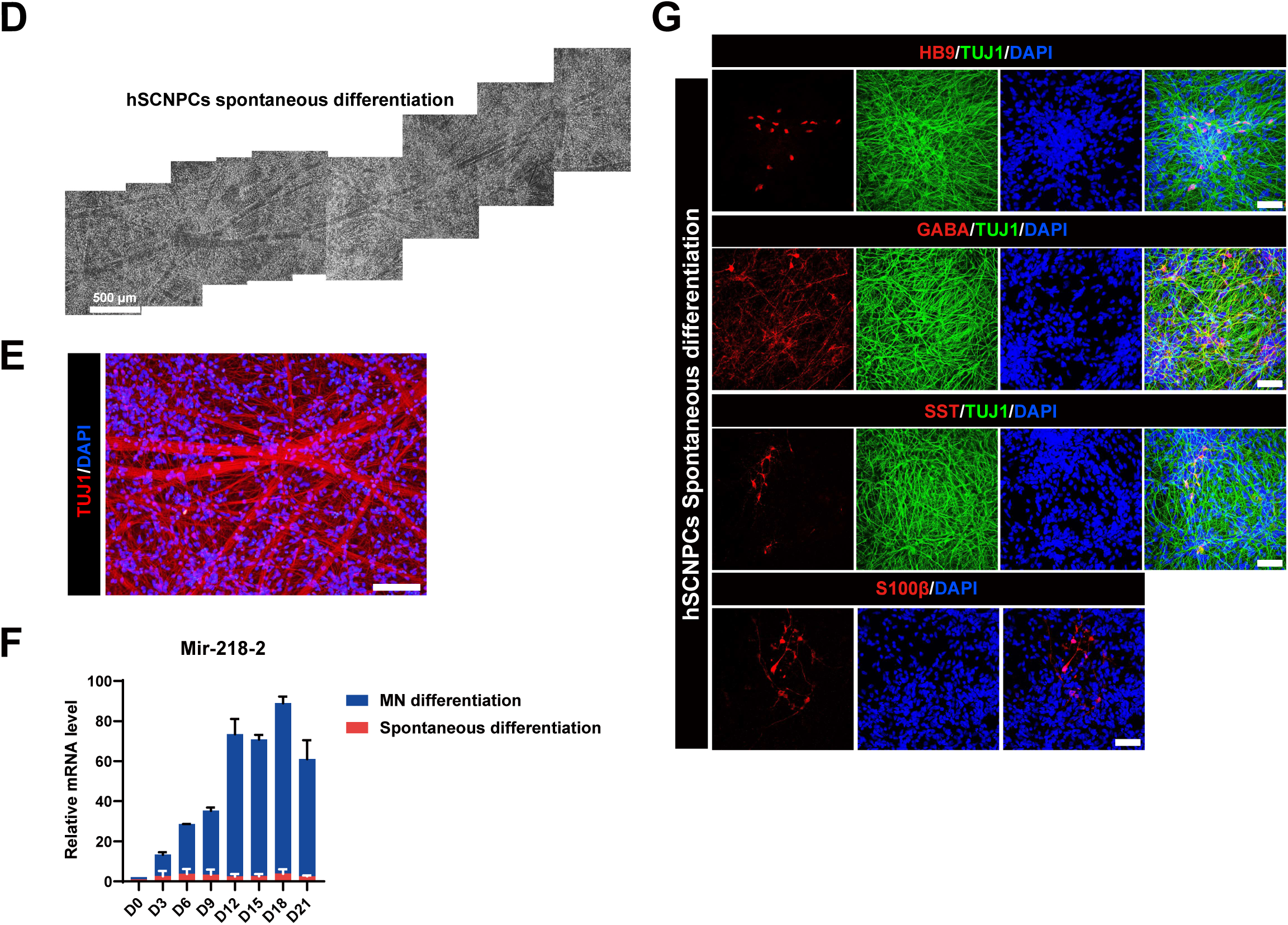
Characterization of hiSCNPCs and multi-potency of hSCNPCs. (A) Relative marker genes expression during spinal motor neuron differentiation from hSCNPCs. n = 3 independent experiments. Data are represented as mean ± SD. (B) Immunostaining characterization of hiSCNPCs (Derived from hiPSCs lines DC60-3 and DC87-3) differentiated spinal motor neurons at day 18. Scale bar, 50 μm. (C) The heat-map showing two regulon groups in samples of spinal motor neuron differentiation from hSCNPCs with listing representative regulon transcription factors (numbers of predicted target genes by SCENIC in the brackets) and enriched GO terms for each regulon group. (D) Bright field of hSCNPCs spontaneous differentiated neurons. Scale bar, 500 μm. (E) Immunofluorescent staining of hSCNPCs spontaneous differentiated neurons with neuronal marker TUJ1. Scale bar, 100 μm. (F) Comparison of mir-218-2 expression levels during hSCNPCs spinal motor neuron differentiation and spontaneous differentiation. n = 2-3 independent experiments. Data are represented as mean ± SD. (G) Immunostaining characterization of hSCNPCs with motor neuron marker (HB9), interneuron marker (GABA, SST) and astrocyte marker (S100β). Scale bar, 50 μm.

**Figure S3.**
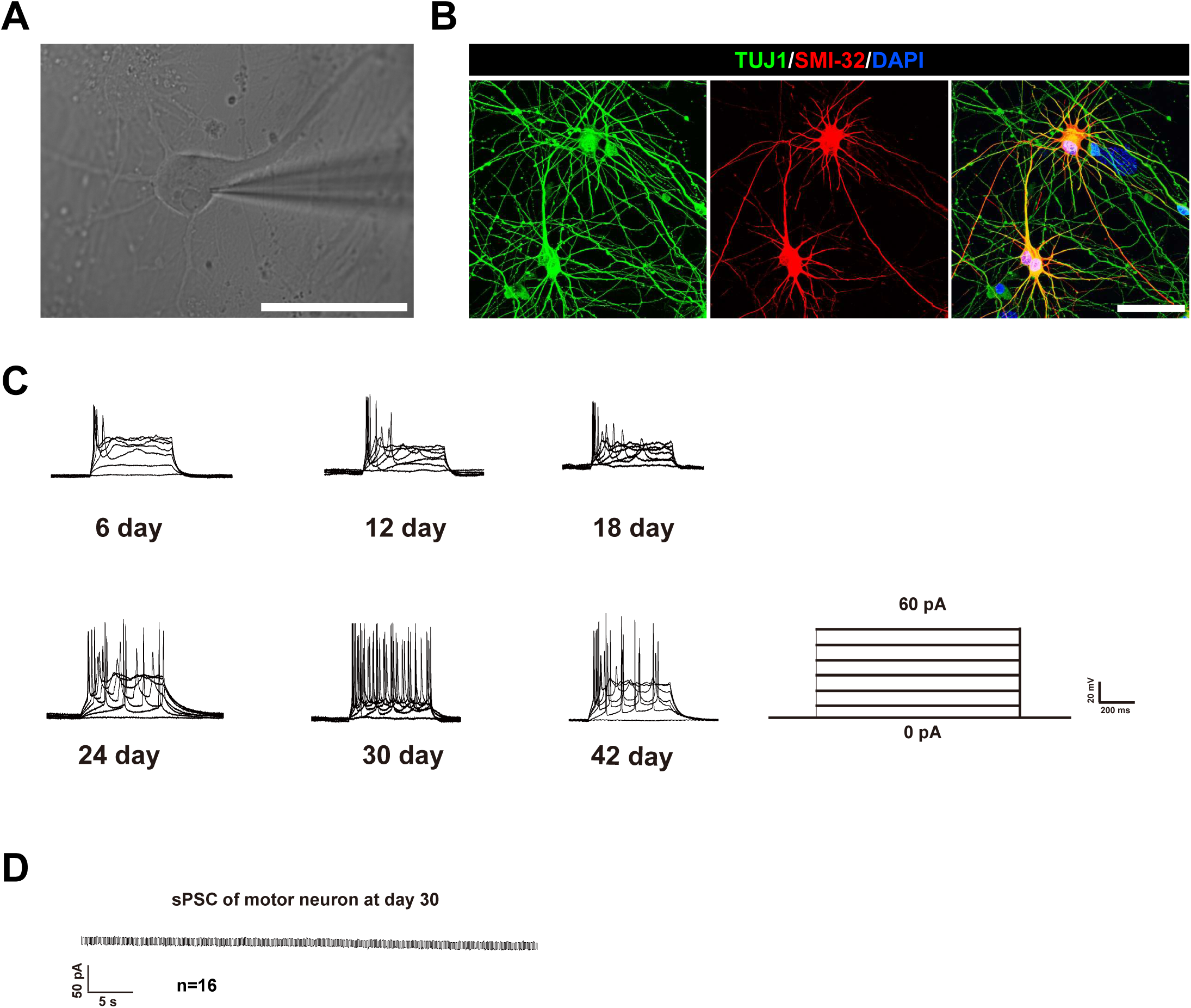
Electrophysiology of hSCNPCs differentiated spinal motor neurons. (A) Bright field illustration of motor neuron under patch clamping. Scale bar, 50 μm. (B) Immunofluorescent staining of TUJ1 and SMI-32 of spinal motor neurons co-culture with astrocyte at day 30. Scale bar, 50 μm. (C) Sample AP traces in response to a series of step current injections from 0 pA to 60 pA. (D) sPSCs of hSCNPCs are not observed in differentiated spinal motor neurons at day 30.

**Figure S4.**
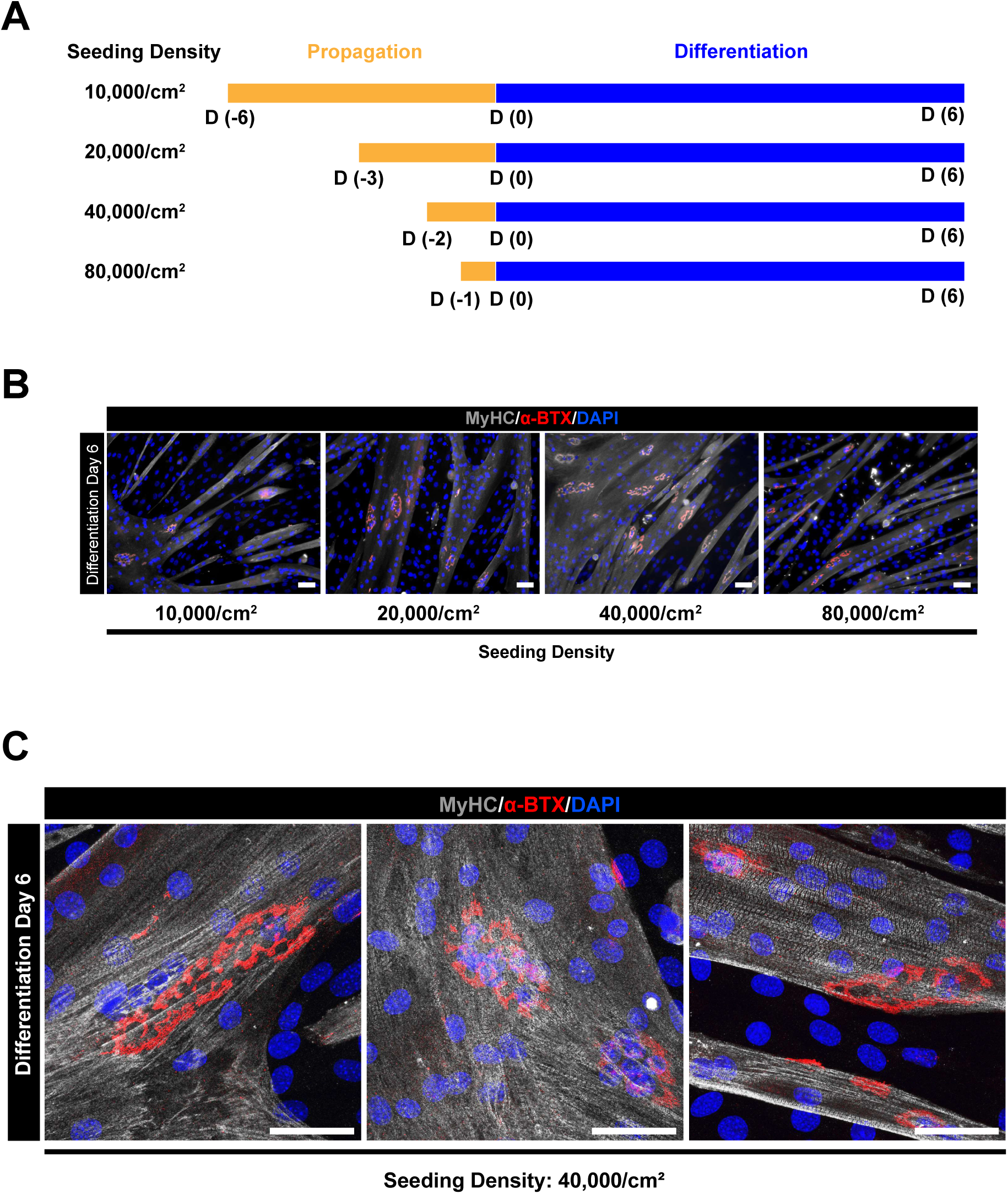
Generation of C2C12 myogenic differentiation. (A) Schematic illustration of C2C12 SOP myogenic differentiation scheme. (B) Immunofluorescent staining of mature muscle marker (MyHC) and acetylcholine receptor accumulation (α-BTX). Scale bar, 200 μm. (C) Representative AChR images at C2C12 differentiated cell at day 6. Scale bar, 50 μm.

