## Supplemental Table S1 for "Generation of functional posterior spinal motor neurons from hPSCs-derived human spinal cord neural progenitor cells"

**Table S1**. Oligonucleotide sequences, related to STAR Methods.

| **RT-qPCR primers** | **Forward** | **Reverse** |
|---|---|---|
| GAPDH | GAGCACAAGAGGAAGAGAGAGACCC | GTTGAGCACAGGGTACTTTATTGATGGTACATG |
| POU5F1 | CGTGAAGCTGGAGAAGGAGAAGCTG | CAAGGGCCGCAGCTTACACATGTTC |
| NKX1-2 | CCCTCCCACCACAAGATTTCT | GACCTCCGCCAAACTTTTCCT |
| CDX2 | GACGTGAGCATGTACCCTAGC | GCGTAGCCATTCCAGTCCT |
| SOX2 | TGGACAGTTACGCGCACAT | CGAGTAGGACATGCTGTAGGT |
| PAX6 | TGGGCAGGTATTACGAGACTG | ACTCCCGCTTATACTGGGCTA |
| HOXB4 | CTGGATGCGCAAAGTTCACGTG | CGTGTCAGGTAGCGGTTGTAGT |
| HOXC9 | CAGCAAGCACAAAGAGGAGAAGG | AGTTCCAGCGTCTGGTACTTGG |
| HOXC10 | GAGCGAAAAGGAGAGGGCCAAA | TCCGCTCTTTGCTGTCAGCCAA |
| NESTIN | GGCGCACCTCAAGATGTCC | CTTGGGGTCCTGAAAGCTG |
| NKX2-2 | CCTTCTACGACAGCAGCGACAA | ACTTGGAGCTTGAGTCCTGAGG |
| NKX6-1 | CCTATTCGTTGGGGATGACAGAG | TCTGTCTCCGAGTCCTGCTTCT |
| HB9 | GATGCCCGACTTCAACTCCC | GCCGCGACAGGTACTTGTT |
| ISL1 | TGAAATGTGCGGAGTGTAATCAGTATTTGGAC | CACACAGCGGAAACACTCGATGTG |
| TUJ1 | GAGCGGATCAGCGTCTACTAC | CCCCACTCTGACCAAAGATGAA |
| NEUN | CCAAGCGGCTACACGTCTC | CGTCCCATTCAGCTTCTCCC |
| VAChT | TTCGCCTCTACAGTCCTGTTC | GCTCCTCCGGGTACTTATCG |
| ChAT | CATGAAGCAATACTATGGGCTCTTCTCCTC | GACGGCGGAAATTAATGACAACATCCAAG |
| Mir-218-2 | TGCGGGGCTTTCCTTTGT | CCGTTTCCATCGTTCCAC |
| gapdh | TGTGATGGGTGTGAACCACGAGAA | CTGTGGTCATGAGCCCTTCCACAA |
| musk | CTGAAGGCTGTGAGTCCACTGT | TCCTTTACCGCCAGGCAGTACT |
| chrng | CTTGTGGCTAAGAAGGTGCCTG | GCAAGGACACATTGAGCACGAC |
| chrne | AGACCTGAGGACACTGTCACCA | TCGTCCTTGCTGTAGTTGAGCC |
